# Alternative transmission patterns in independently acquired nutritional co-symbionts of Dictyopharidae planthoppers

**DOI:** 10.1101/2021.04.07.438848

**Authors:** Anna Michalik, Diego C. Franco, Michał Kobiałka, Teresa Szklarzewicz, Adam Stroiński, Piotr Łukasik

**Affiliations:** Department of Developmental Biology and Morphology of Invertebrates, Institute of Zoology and Biomedical Research, Faculty of Biology, Jagiellonian University, Gronostajowa 9, 30-387 Kraków, Poland; Institute of Environmental Sciences, Faculty of Biology, Jagiellonian University, Gronostajowa 7, 30-387 Kraków, Poland; Museum and Institute of Zoology, Polish Academy of Sciences, Wilcza 64, 00-679 Warszawa, Poland

**Author notes:** corresponding authors: Anna Michalik.

**Keywords:** planthoppers, nutritional endosymbiosis, transovarial transmission

## Abstract

Sap-sucking hemipterans host specialized, heritable microorganisms that supplement their unbalanced diet with essential nutrients. These microbes show unusual features that provide a unique perspective on the evolution of life but have not been systematically studied. Here, we combine microscopy with high-throughput sequencing to revisit 80-year-old reports on the diversity of symbiont transmission modes in a broadly distributed planthopper family Dictyopharidae. We show that in all species examined, the ancestral nutritional endosymbionts *Sulcia* and *Vidania* are complemented by co-primary symbionts, either *Arsenophonus* or *Sodalis*, acquired several times independently by different host lineages. Like in other obligate sap-feeders, the ancestral symbionts produce essential amino acids, whereas co-primary symbionts contribute to the biosynthesis of B vitamins. These symbionts reside within separate bacteriomes within the abdominal cavity, although in females, *Vidania* also occupies bacteriocytes in the rectal organ. Notably, the symbionts are transmitted from mothers to offspring in two alternative ways. In most examined species, all nutritional symbionts simultaneously infect the posterior end of the full-grown (vitellogenic) oocytes and next gather in their perivitelline space. In contrast, in other species, *Sodalis* colonizes the cytoplasm of the anterior pole of young (previtellogenic) oocytes forming a cluster separate from the “symbiont ball” formed by late-invading *Sulcia* and *Vidania*. Our data add to evidence on frequent replacements of gammaproteobacterial symbionts combined with the relative functional stability of the nutritional functions during the evolution of sap-feeding insects, and show how newly-arriving microbes may utilize different strategies to establish long-term heritable symbiosis.

**Significance statement:** Sup-sucking hemipterans host ancient heritable microorganisms that supplement their unbalanced diet with essential nutrients, and which have repeatedly been complemented or replaced by other microorganisms. They need to be reliably transmitted to subsequent generations through the reproductive system, and often they end up using the same route as the ancient symbionts. We show for the first time that in a single family of planthoppers, the complementing symbionts that have established infections independently utilize different transmission strategies, one of them novel, with the transmission of different microbes separated spatially and temporarily. These data show how newly-arriving microbes may utilize different strategies to establish long-term heritable symbiosis.

## Introduction

Mutualistic relationships with heritable bacterial and/or fungal microorganisms have played crucial roles in the biology of multiple groups of insects, contributing significantly to their evolutionary and ecological success (1–3). The growing awareness of the diversity and importance of insect symbioses, in addition to the rapid development in sequencing-based techniques, has led to an increased interest in these associations. However, outside a few model species and some reasonably well-sampled clades, our knowledge of the diversity, evolution, and biological characteristics of the microbial symbionts, and the microbial roles in the evolution of insect diversity remains limited (4, 5). Among insects, sap-sucking hemipterans are obligatorily dependent on heritable nutritional microbes that supplement their unbalanced diet with essential amino acids, vitamins, and co-factors (4, 6–8). Multiple symbiont combinations have been described from Auchenorrhyncha, a suborder comprising infraorders Fulgoromorpha – planthoppers, and Cicadomorpha – cicadas, spittlebugs, treehoppers, and leafhoppers (4, 9). Their common ancestor that lived about 300 million years ago (MYA) is thought to have became colonized by two microbes, a Bacteroidetes currently known as *Candidatus* Sulcia muelleri (further referred to as *Sulcia*) and a beta proteobacterium, variably known as *Candidatus* Nasuia deltocephalinicola, *Candidatus* Zinderia insecticola or *Candidatus* Vidania fulgoroidaea (further referred to as *Nasuia*, *Zinderia*, and *Vidania,* respectively, or as beta-symbionts collectively) (9–12). These nutrient-providing symbionts have become obligate components of the host biology, and transmitting strictly maternally, they have co-diversified with hosts. However, in many host clades, one or both became complemented or replaced by other microbes. For example, an Alphaproteobacterium *Candidatus* Hodgkinia cicadicola (further referred to as *Hodgkinia*) has replaced the beta-symbiont in the ancestor of modern cicadas (family Cicadidae), but has itself became repeatedly replaced by fungi from the genus *Ophiocordyceps* (13, 14).

In these multi-partite symbioses, microbes share the responsibility for essential nutrient biosynthesis. For example, in known Cicadomorpha, *Sulcia* encodes pathways for producing 7 or 8 essential amino acids, whereas the remaining 3 or 2 amino acids are provided by their symbiotic partner (8, 9, 12). The situation can get more complicated when one of the ancient microbes gets replaced by another, or when the host is colonized by more than two symbionts and the nutritional functions become subdivided among a greater number of partners. This has occurred repeatedly in different lineages of Auchenorrhyncha, where additional symbionts have either taken over beta-symbiont’s role or contribute to vitamin biosynthesis (4).

These intimate and intricate metabolic interdependencies between insects and associated microorganisms indicate a vital role of nutritional symbionts in host biology, and the need to ensure reliable transmission of these essential partners across generations. In sap-feeding hemipterans, transovarial transmission through female germ cells predominates (15–17). It may take place at all stages of female germ cell development. In some insects, symbionts infect undifferentiated germ cells, or alternatively, young, previtellogenic oocytes (17). However, in most hemipteran taxa, including all Auchenorrhyncha studied to date, they invade ovarioles containing older (vitellogenic or choriogenic) oocytes. The complementation or replacement of ancient, co-adapted heritable nutritional symbionts by newly arriving microbes, while likely beneficial to the hosts because of their greater metabolic capacity and efficiency, creates apparent challenges for their transmission. Host lineages that acquired new symbionts have to develop new traits and mechanisms of their effective vertical transmission, which is crucial for the fixation of new symbiosis (18), being also a matter of life or death to the newly established symbionts. The evolution of the symbiont transmission and the symbiont replacements are inseparably linked, and we need to study one to understand another.

Buchner (15) while summarizing decades of microscopy-based research on Auchenorrhyncha symbioses, famously wrote about “the veritable fairyland of insect symbiosis”, apparently referring to the diversity of microbes he observed in different host clades, as well as their transmission mechanisms. However, he lacked tools to fully characterize the evolution of symbioses across the auchenorrhynchan phylogeny. The popularization of DNA sequencing-based techniques enabled such investigation, but surprisingly, our knowledge of these links remains limited to a few Auchenorrhyncha clades, mainly within Cicadomorpha (12, 18, 19). While diagnostic screens revealed *Sulcia*, *Vidania*, and often other bacteria or fungi in most planthopper families, in only one planthopper species so far nutritional symbionts have been characterized using genomics. In the Hawaiian cixiid *Oliarus filicicola*, *Vidania* produces seven essential amino acids, *Sulcia* synthesizes three, whereas a more recently acquired gammaproteobacterial symbiont *Purcelliella* contributes B-vitamins. This rearrangement of *Sulcia* and *Vidania* nutritional responsibilities shed new light on the evolution of planthopper symbioses and how they can be influenced by infections with additional microbes. However our understanding of symbioses in this diverse, widespread, and ecologically significant insect clade remains very limited.

This project aimed to provide an insight into the diversity and biology of symbioses in the large, diverse planthopper family Dictyopharidae. Eight members of this family were shown to possess bacteria *Vidania* and usually also *Sulcia* (20), but microscopic observations conducted by Müller (21, 22) and summarized by Buchner (15) indicated that Dictyopharidae also host the third symbiont.

Strikingly, they reported that in different Dictyopharidae species, these additional symbionts can be transmitted in alternative ways, either together or separately from *Sulcia* and *Vidania.* While this suggested the independent origins of the additional symbionts, or other unusual biological or evolutionary processes, Buchner and colleagues lacked the tools to fully describe and explain the phenomenon. There was, however, little doubt that Dictyopharidae can provide unique insights into the symbiont evolution, transmission, and replacements, making this family a valuable model group for the exploration of symbiont complementation and genome evolution.

Here, we report the results of microscopy and sequencing-based investigations of symbiotic bacteria associated with seven species belonging to the Dictyopharidae. We survey the diversity of the symbionts and explore their nutritional roles. In particular, we focus on how symbionts in this group are transmitted across generations. We describe and discuss how the auchenorrhynchan symbiont transmission can be separated in time and space, how this may have evolved, and what it may mean for the host.

## Results

### Dictyopharidae planthoppers harbor (at least) three types of nutritional symbionts

Analyzes of marker gene sequences for the representative specimens of seven experimental species confirmed their morphology-based identifications (Fig. 1A, *SI Appendix*, Figs. S3, and S4). The phylogeny revealed two well-supported clades corresponding to subfamilies Dictyopharinae and Orgerinae; it also showed that *R. edirneus* is more closely related to *P. platypus* than to *R. scytha*, in agreement with the proposed taxonomic revision of the group (Emeljanow, 2003). We successfully amplified bacterial 16S rRNA gene V4 region from the abdomens of all experimental individuals from these seven Dictyopharidae species. The total number of 16S rRNA reads passed through all analysis steps was 452,949, or 23,840 per sample on average (Dataset Table S5). Clustering with 97% identity cutoff identified 67 OTUs.

**Figure 1.**
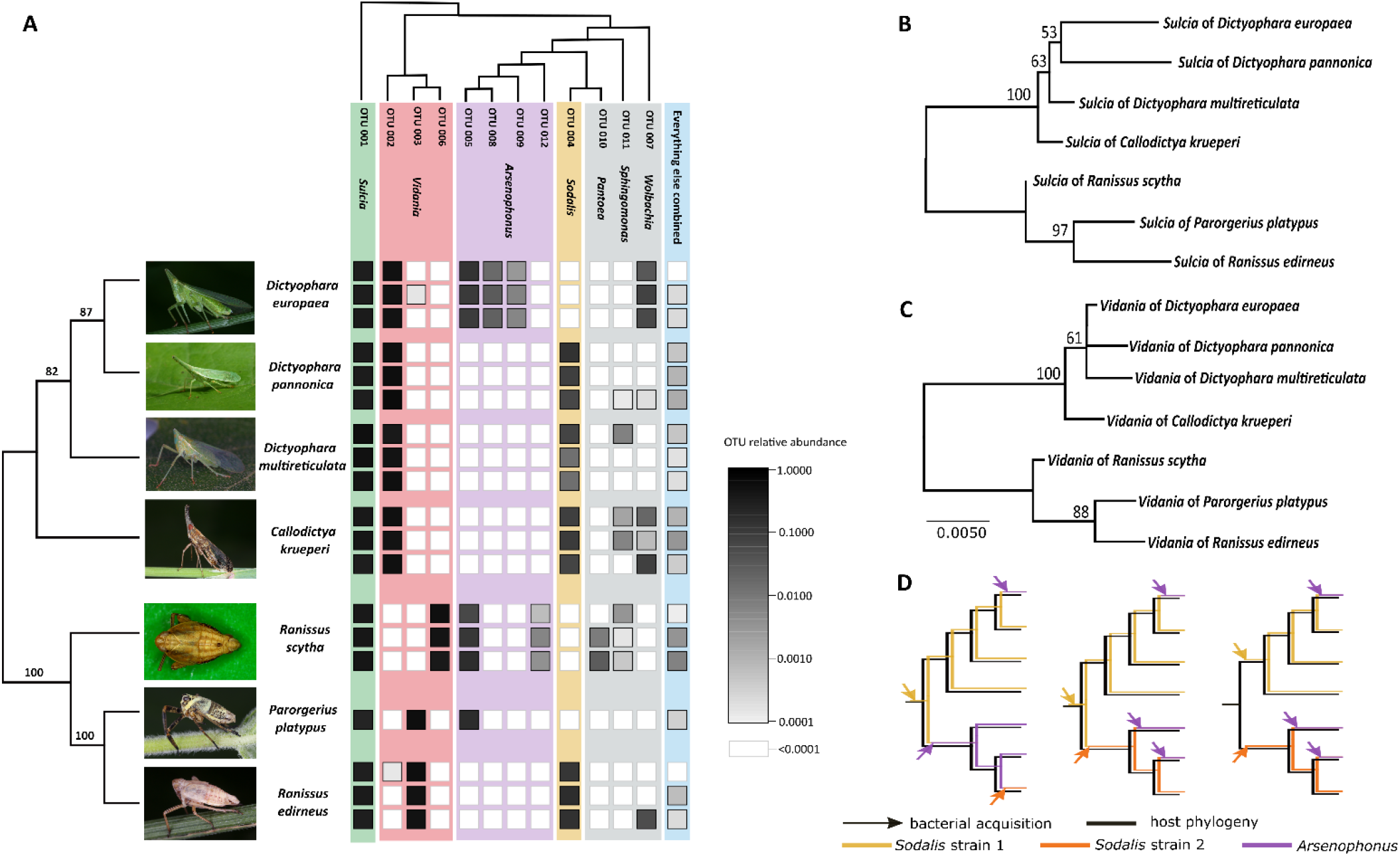
The bacterial communities in Dictyopharidae planthoppers. **A.** The diversity and relative abundance of bacteria in replicate specimens from seven experimental species. Insect phylogeny (Maximum Likelihood) is based on concatenated marker gene sequences (COI, CytB, 18S rRNA, and 28S rRNA); bootstrap values are shown above the nodes. The bacterial tree is based on representative sequences of 16S rRNA V4 region for different OTUs. **B & C.** ML phylogenies for *Sulcia* and *Vidania* symbionts from seven experimental species, based on full-length 16S rRNA sequences. **D.** Three of many possible scenarios of *Sodalis*/ *Arsenophonus* acquisition and replacement during Dictyopharidae diversification.

All studied individuals hosted *Sulcia* and *Vidania*, and additionally, either *Sodalis* or *Arsenophonus* (Fig. 1A, *SI Appendix*, Fig. S1, Dataset Table S5). Together, these symbionts comprised >97% of reads in each of the libraries. The single-nucleotide-resolution data (Dataset Table S6) for these dominant symbionts revealed that all species harbor very similar strains of the slow-evolving symbiont *Sulcia*, all clustering to a single 97% OTU. In contrast, *Vidania* genotypes from different species were sufficiently divergent to fall into three distinct 97% OTUs. *C. krueperi*, *D. multireticulata*, *D. pannonica*, and *R. edirneus* hosted different genotypes of *Sodalis* (clustering to the same OTU).*D. europaea*, *S. scytha* and *P. platypus* hosted *Arsenophonus*, typically different genotypes from more than one OTU. However, their consistent relative abundance in replicate individuals strongly suggests that these genotypes and OTUs correspond to different rRNA operons within the genome of a single *Arsenophonus* strain (Fig. 1A). Less abundant microbial OTUs present in some species included *Wolbachia*, *Pantoea*, and *Sphingomonas* (Fig. 1A). All the remaining OTUs combined accounted for 0.2% of the total number of reads, and many of them represented contaminants or symbiont-derived sequences that accumulated large numbers of errors. We did not consider them further.

Phylogenetic trees of *Sulcia* and *Vidania* based on full-length 16S rRNA gene sequences (Fig. 1B, C) are congruent with the phylogeny of the seven experimental Dictyopharidae species (Fig. 1A), or a broader range of planthoppers (Suppl. Figs S2 and S3), as expected for symbionts co-diversifying with hosts. In contrast, 16S rRNA gene phylogenies for *Sodalis* and *Arsenophonus* from diverse hosts, while poorly resolved and supported, were suggestive of independent origins of at least some of co-symbionts of Dictyopharidae (*SI Appendix*, Figs. S4 and S5). These data, combined with information on the distribution of the two symbiont clades across the host phylogeny and differences in transmission patterns and genomics characteristics (described later), are suggestive of several independent infections by *Arsenophonus*/*Sodalis*, or their repeated replacements by other strains of these symbionts (Fig 1B, C, D, *SI Appendix*, Figs 2-5). Unfortunately, the currently available data do not allow for a reconstruction of the order of these infections; we lack the resolution to distinguish among the many possible scenarios.

**Figure 2.**
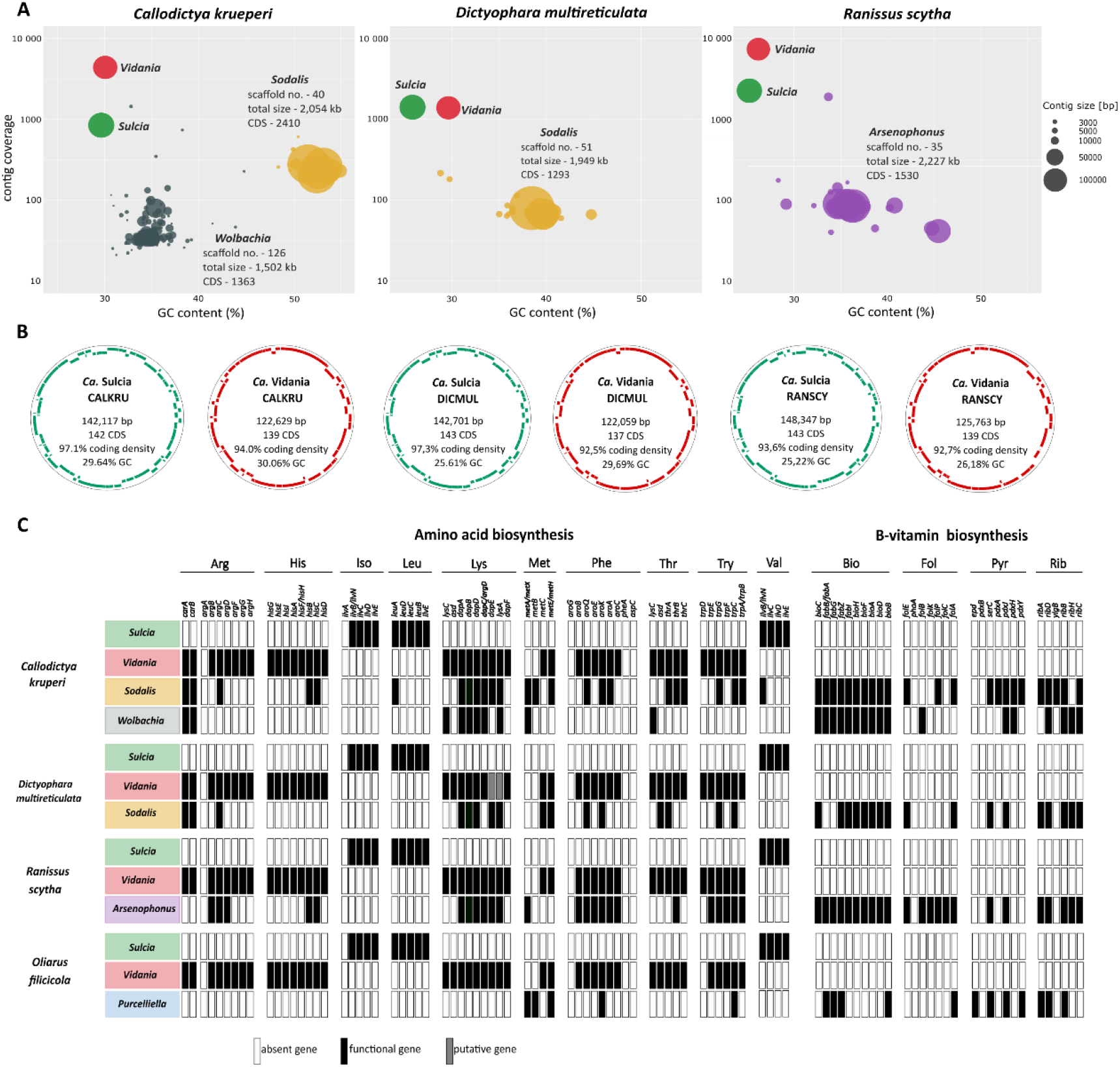
The summary of metagenomic data for three Dictyopharidae species. **A.** Taxon-annotated GC contents-coverage plots for metagenomic assemblies. Colored blobs represent scaffolds corresponding to the identified symbiont genome fragments. **B.** Circular diagrams of *Sulcia* and *Vidania* genomes from three species, showing gene positions on forward and reverse strands, and basic genome characteristics. **C.** Contribution of symbionts to amino acid and B-vitamin biosynthesis pathways. Standard abbreviations for amino acid, B-vitamin, and gene names are used.

Three of these scenarios, all assuming relatively few replacement events and shared ancestry of some strains, are presented in Fig. 1D. However, the number of independent infections or replacements along the tree branches could have been much greater. In fact, patterns such as the 16S rRNA amplicon genotype diversity across *D. pannonica* individuals (Dataset Table S7) and some morphological differences among *Sodalis* cell populations hint at the possibility of recent or perhaps ongoing replacements.

### *Sulcia* and *Vidania* provide essential amino acids to host insects, whereas *Arsenophonus*, *Sodalis*, and *Wolbachia* supplement their diet with B - vitamins

The symbiont genomes identified in the assemblies matched the dominant OTUs identified by amplicon sequencing (Fig. 2A). All species hosted *Sulcia* and *Vidania*, and in all cases, we were able to assemble circular genomes with sizes ranging between 142-148 kb for *Sulcia* and 122-125 kb for *Vidania,* characterized by relatively low GC contents (25-30%) and high coverage (800-10000x) (Fig. 2B). *Sulcia* and *Vidania* genomes were co-linear relative to each other and the slightly larger reference genomes of *O. filicicola* (OLIH) symbionts (157 and 136 kb, respectively - Bennett & Mao 2018; Suppl. Figs S6 and S7). Our metagenomic analyses also confirmed the presence of *Sodalis* and *Wolbachia* in CALKRU, *Sodalis* in DICMUL, and *Arsenophonus* in RANSCY. The genomic assemblies of these symbionts were fragmented; the number of scaffolds ranged between 23-126 and their total size between 1500 kb and 2220 kb. The scaffold read coverage was substantially lower than in the case of *Sulcia*/*Vidania* but also relatively variable, particularly in the case of *Arsenophonus* (Dataset Table S7).

In the terms of gene contents, the newly characterized *Sulcia* and *Vidania* genomes are very similar to each other and the previously described OLIH symbiont genomes (Bennett and Mao, 2018). *Sulcia* genomes encode 142-143 predicted protein-coding genes, 27-29 identifiable tRNAs, complete ribosomal operon, and their coding density ranges from 93.6% to 97.3%. *Vidania* genomes encode 137-139 predicted protein-coding genes, 23-25 tRNAs, complete ribosomal operon and coding density ranging between 92.5%-94% (Fig. 2B). In the two analyzed *Sodalis* genomes, prokka identified 2410 (CALKRU) and 1132 (DICMUL) predicted protein-coding genes, in *Arsenophonus* 1530, and in *Wolbachia*, 1363. However, because of the incompleteness of the assemblies of these co-symbiont genomes and challenges with pseudogene annotation, these numbers are approximate.

Our analyses of ancestral symbionts *Sulcia* and *Vidania* protein-coding genes revealed that they complement each other to supplement host insects with essential amino acids, in a way consistent among the three Dictyopharidae and OLIH (Fig. 2C). *Sulcia* participates in the biosynthesis of three out of ten essential amino acids, including isoleucine, leucine, and valine, whereas *Vidania* is involved in synthesizing the remaining seven. Interestingly, some of the genes in amino acid biosynthetic pathways (isoleucine, arginine, methionine, phenylalanine) were not found in *Sulcia* and *Vidania* genomes (Fig. 2C). Some of these missing amino acid biosynthesis genes, and duplicate copies of some others, are present in *Sodalis, Arsenophonus*, and *Wolbachia* genomes. In *C. krueperi*, *Vidania* lacks the first two genes essential for methionine biosynthesis from homoserine; however, these missing genes are present in the *Sodalis* genome. Similarly, in *D. multireticulata*, *Sodalis* complements two pseudogenes (*dapE*, *lysA*) in the *Vidania* genome essential in the lysine biosynthesis pathway. *Arsenophonus* from *R. scytha* has the same set of genes from phenylalanine and tryptophan biosynthesis as *Vidania*. In all Dictyopharidae, the gammaproteobacterial symbionts encode many genes in the lysine biosynthesis pathway and scattered genes from other pathways. We can not rule out the presence in the third co-primary symbiont genomes’ remaining genes missing in *Sulcia* and *Vidania* due to the uncompleted assembly process.

In addition to genes that might participate in amino acids’ biosynthesis, *Sodalis, Arsenophonus*, and *Wolbachia* contribute to the synthesis of B-vitamins: biotin, folate, riboflavin, and pyridoxine (Fig. 2C). *Sodalis* and *Wolbachia* associated with *C. krueperi* as well as *Arsenophonus* of *R. scytha* encode complete sets of biotin biosynthesis genes, whereas the genome of *Sodalis* of *D. multireticulata* carries only 8 of 10 genes from that pathway. *Arsenophonus* symbiont genome also encodes almost all genes involved in the biosynthesis of folate and riboflavin. Interestingly, in *C. krueperi,* the riboflavin pathway is shared by *Sodalis* and *Wolbachia*.

### Nutritional symbionts of dictyopharids occupy distinct but adjacent bacteriocytes

In all dictyopharids studied, we found nutritional symbionts (*Sulcia*, *Vidania*, and either *Sodalis* or *Arsenophonus*) contained within separate bacteriomes located in proximity to each other within the insect abdomens (Figs 3, 4). Light microscopy observations have revealed that bacteriomes harboring bacteria *Sulcia*, *Sodalis*, and *Arsenophonus* are made up of several bacteriocytes (Figs 3; 4A), whereas bacteriomes with *Vidania* are syncytial (Figs 3, 4D). Bacteriomes occupied by *Sulcia* and *Vidania* are surrounded by thick (in *Sulcia*) or thin (in *Vidania*) monolayered bacteriome sheath (Fig. 4A, C, D, F). In contrast, bacteriomes with *Arsenophonus* and *Sodalis* are not covered by epithelial cells (Fig. 3).

**Figure 3.**
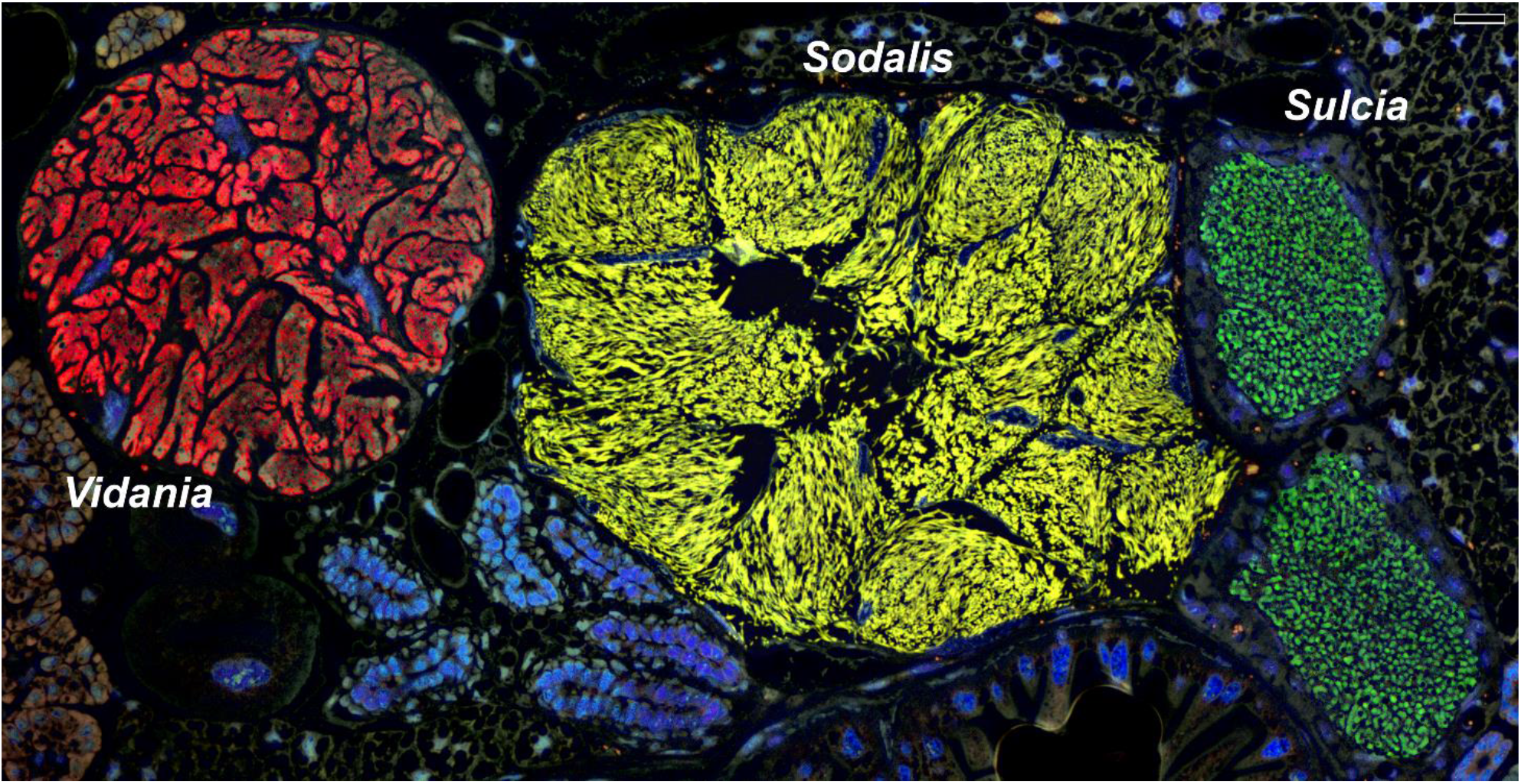
Fluorescent *in-situ* hybridization (FISH) demonstrates how in *C. krueperi*, each of its three co-primary symbionts inhabit distinct bacteriomes. Specific probes for *Vidania* (red), *Sulcia* (green), and *Sodalis* (yellow) were used. Blue represents DAPI. Confocal laser microscope (CLM), scale bar = 20 μm.

**Figure 4.**
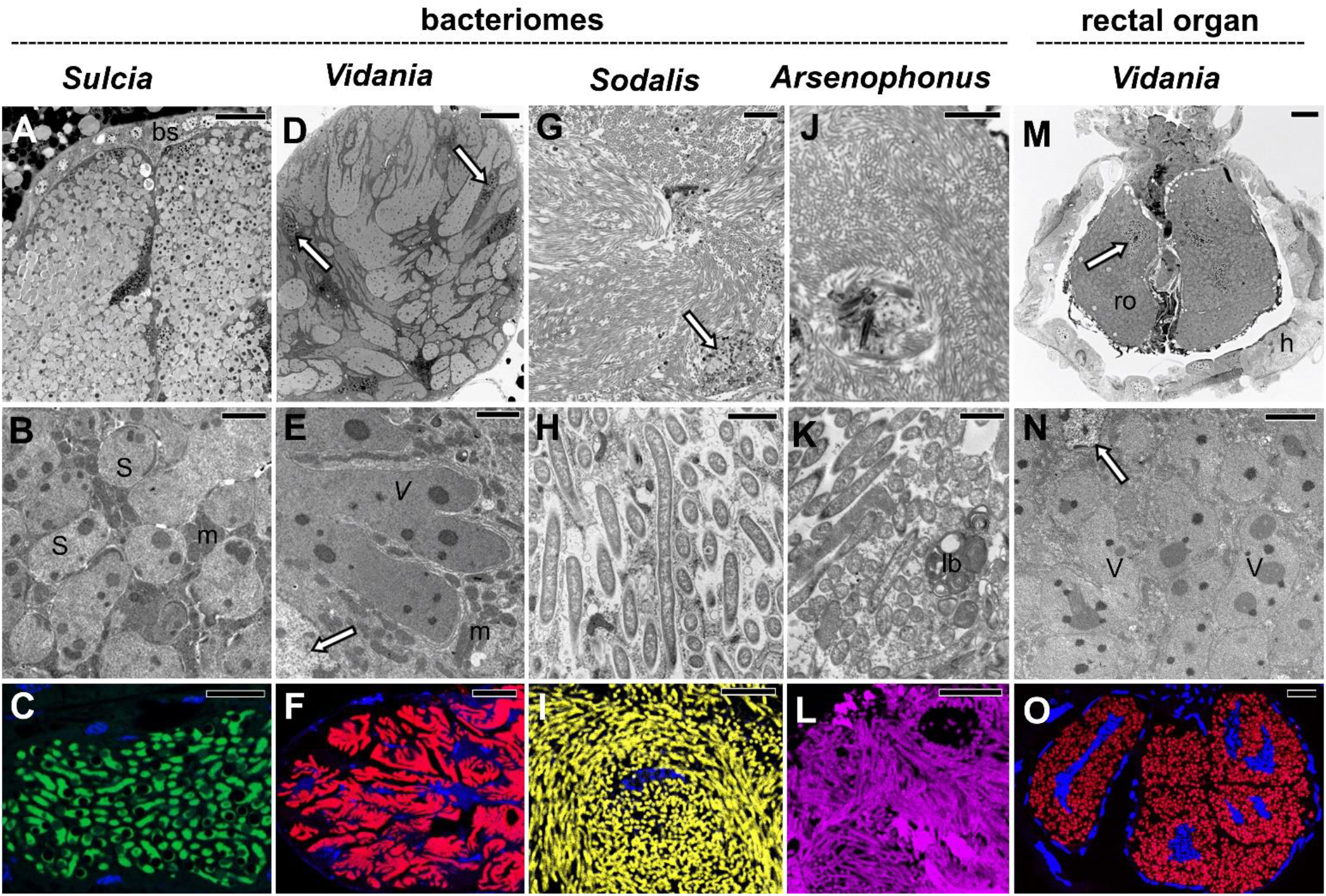
Tissue localization and morphology of symbionts in the Dictyopharidae species examined. **Top row (A, D, G, J, M):** the organization of symbionts within bacteriomes or the rectal organ. Light microscopy (LM), scale bar = 20 μm. **Middle row (B, E, H, K, N):** the ultrastructure of symbiont cells. Transmission electron microscopy (TEM), scale bar = 2μm. **Bottom row (C, F, I, L, O):** fluorescent *in-situ* hybridization (FISH) microphotographs of symbiont cells within the bacteriome or rectal organ. Probes specific to each of the symbionts were used. Blue represents DAPI. Confocal laser microscope (CLM), scale bar = 20 μm. Insect species: **A -** *D. multireticulata*, **B, L -** *R. scytha*, **C, E-I, M, N -** *C. krueperi*, **D, O -** *D. pannonica*, **J, K -** *D. europaea*. Abbreviations: bs – bacteriome sheath, h – hindgut, lb – lamellar body, m – mitochondrion, ro – rectal organ, S – *Sulcia*, V – *Vidania*, white arrow – bacteriocyte nucleus.

Our histological, ultrastructural, and FISH analyses revealed that *Sulcia* cells are pleomorphic and possess large electron-dense inclusions in their cytoplasm (Figs. 3; 4A-C). *Vidania* cells are giant and multi-lobed, with numerous electron-dense accumulations in the cytoplasm (Figs. 3; 4D-F). The third type of nutritional symbionts - bacteria *Sodalis* (in *C. krueperi*, *D. pannonica*, *D. multireticulata*, and *R. edirneus*) and *Arsenophonus* (in *D. europaea*, *R. scytha*, and *P. platypus*) are large and elongated (Figs 3, 4J-L). As a rule, their morphology is very similar within species. In particular, morphological data did not suggest the presence of distinct *Arsenophonus* strains, further supporting our amplicon-based conclusions that different 16S OTUs correspond to different rRNA operons of the same strains (Figs 3; 4J-L). The only exception was *Sodalis* in *D. pannonica*, where in some individuals, we observed two different morphotypes of these bacteria, differing in size, methylene blue staining intensity, and electron density. These morphotypes share the same bacteriocytes; however, they may be mixed or form aggregates (not shown).

In all species analyzed, bacteriocytes have large, polyploid nuclei, and their cytoplasm is tightly packed with symbionts, ribosomes, and mitochondria (Figs. 3; 4). The density of mitochondria in bacteriocytes with *Sulcia* and *Vidania* seemed substantially higher than in bacteriocytes with *Sodalis* and *Arsenophonus*. Bacteria *Sulcia* and *Vidania* tightly adhere to each other in the cytoplasm of their bacteriocytes, whereas *Sodalis* and *Arsenophonus* are somewhat less densely packed (Figs 3; 4A-L). Finally, during ultrastructural observations in *C. krueperi* and *D. europaea*, we detected small, rod-shaped bacteria in the cytoplasm of bacteriocytes housed by *Sulcia* and *Vidania* symbionts (*SI Appendix*, Fig. S9). Their shape and size matched *Wolbachia*, detected in these species using amplicon sequencing.

Besides bacteriomes, *Vidania* also occupies bacteriocytes in the rectal organ, localized in the invagination of hindgut epithelium (Fig. 4M-O). Both histological and ultrastructural studies made it clear that *Vidania* cells in the rectal organ differ in shape and size from *Vidania* localized in the bacteriomes.

### Dictyopharids developed different modes of symbiont transmission

Histological observations of serial semithin sections have shown that all nutritional symbionts detected in Dictyopharidae species analyzed are transovarially transmitted between generations. The details of *Sulcia*, *Vidania,* and *Arsenophonus* transmission agree with observations from other Auchenorrhyncha. Strikingly, we observed significant differences in the modes of transmission of *Sodalis* (Figs 5-8).

**Figure 5.**
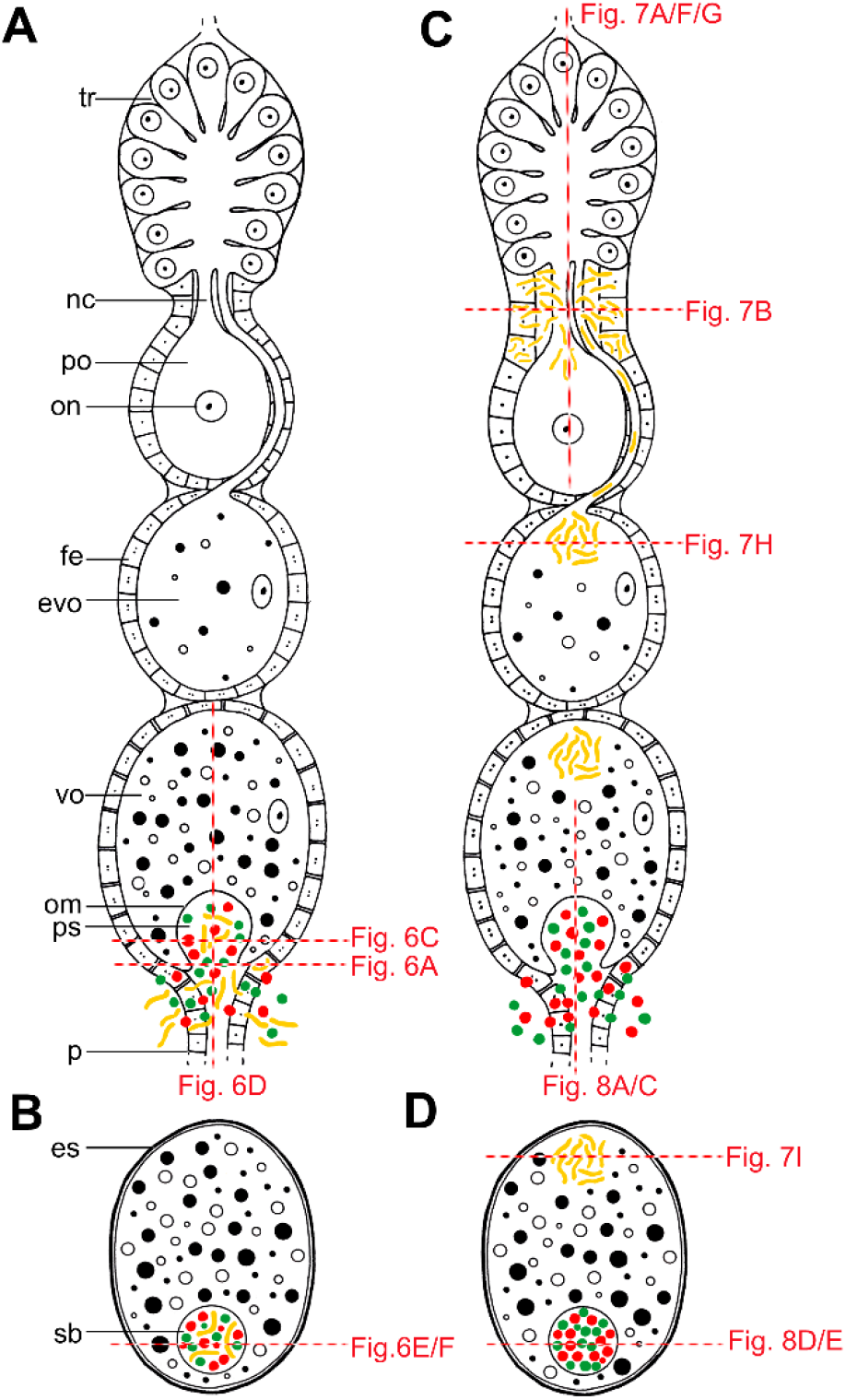
Schematic representation of the alternative modes of the symbiont transmission in Dictyopharidae planthoppers. **A.** Simultaneous transmission of all types of nutritional symbionts through the follicular epithelium surrounding the posterior pole of terminal oocyte. **B.** Full-grown oocyte with a symbiont ball containing three types of symbionts. The transmission mode shown in A-B is used by the large majority of Auchenorrhyncha, including subfamily Orgerinae and *Arsenophonus*-infected *D. europaea* from subfamily Dictyopharinae. **C.** Spatially and temporarily separated transmission of different symbionts (*Sodalis* vs. *Sulcia* and *Vidania*). *Sodalis* infects previtellogenic and early-vitellogenic oocytes, whereas the remaining symbionts invade terminal vitellogenic oocytes. **D.** Full-grown oocyte with the accumulation of *Sodalis* bacteria at the anterior pole, and symbiont ball with *Sulcia* and *Vidania* cells at the posterior pole. This mode of transmission is unique to *Sodalis*-infected members of the subfamily Dictyopharinae. Red dashed lines indicate regions shown on panels of Figures 6-8. Abbreviations: tr – tropharium, nc – nutritive cord, po – previtellogenic oocyte, on – oocyte nucleus, fe – follicular epithelium, evo – early-vitellogenic oocyte, vo – vitellogenic oocyte, om – oocyte membrane, ps – perivitelline space, es – egg shells, sb – symbiont ball.

In all species examined, *Sulcia* and *Vidania* infect the posterior end of the ovariole, migrating at the same time (Figs 5A; 6A).

In contrast, in *C. krueperi*, *D. pannonica*, and *D. multireticulata*, we observed the separation of symbiont transmission in time and space. Instead of entering the perivitelline space through follicular cells, *Sodalis* infects the neck region of the ovariole i.e. the part of the ovariole between tropharium and vitellarium. We observed large accumulations of *Sodalis* cells in ovarioles containing oocytes in the previtellogenic stage of oogenesis. The follicular cells separating the tropharium from the vitellarium are tightly packed with bacteria (Fig 7A-C). This specific region of the ovariole is penetrated by nutritive cords which connect oocytes developing in the vitellarium with trophocytes localized in the tropharium (Fig. 7A-C). Unfortunately, due to the lack of larval stage of species analyzed, we could not observe the migration of *Sodalis* symbionts from the body cavity to the ovariole. Bacteria *Sodalis* infect previtellogenic oocytes using nutritive cords (Fig. 7D-H). They leave the follicular cells *en masse*, enter the nutritive cord area (Fig. 7D, E) and migrate towards the oocyte along microtubules (Fig. 7F-H). Then, *Sodalis* cells aggregate in the cytoplasm of the anterior pole of the oocyte (Fig. 7H) and stay in this form through the next stages of oogenesis (Fig. 7I). In these species, *Sulcia* and *Vidania* symbionts are transmitted to the perivitelline space of choriogenic oocytes (which contain clusters of *Sodalis* bacteria in the cytoplasm of the anterior pole) by way of the follicular cells at their posterior end (Fig. 8A-C). They closely adhere to each other and form a characteristic “symbiont ball” (Fig. 8D-F).

**Figure 6.**
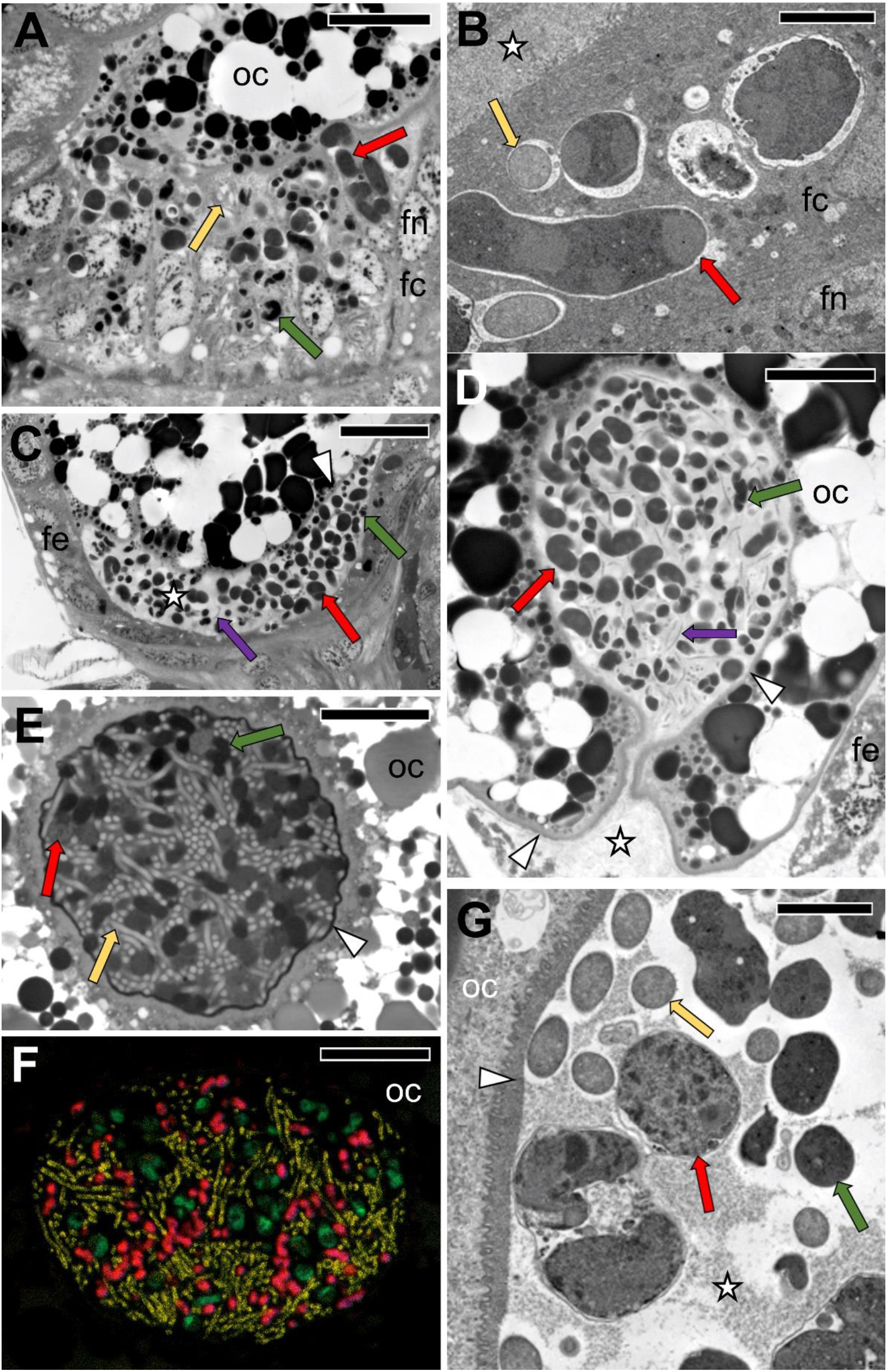
The simultaneous transmission of three symbionts in selected Dictyopharidae, as shown in Fig. 5A-B. **A.** The migration of *Sulcia*, *Vidania,* and *Arsenophonus* through follicular cells surrounding the posterior pole of the terminal oocyte. *D. europaea,* light microscopy (LM), scale bar = 20 μm. **B.** *Sodalis* and *Vidania* in the cytoplasm of the follicular cell. *R. edirneus,* TEM, scale bar = 2 μm. **C, D.** Symbiotic bacteria in the perivitelline space. *D. europaea*, LM, scale bar = 20 μm. **E.** A ‘symbiont ball’ containing bacteria *Sulcia*, *Vidania,* and *Sodalis* in the deep depression of the oolemma at the posterior pole of the terminal oocyte. *R. edirneus*, LM, scale bar = 20 μm. **F.** *In situ* identification of symbionts in the “symbiont ball” within the terminal oocyte. *R. edirneus*, CLM, scale bar = 20 μm. **G.** Fragment of the “symbiont ball” in the perivitelline space. *R, edirneus*, TEM, scale bar = 2 μm. Abbreviations: fc – follicular cell, fn – the nucleus of the follicular cell, oc – oocyte, asterisk – perivitelline space, arrowhead – oocyte membrane, red cells/arrows – *Vidania*, green cells/arrows – *Sulcia*, yellow cells/arrows – *Sodalis*, purple arrows – *Arsenophonus*

**Figure 7.**
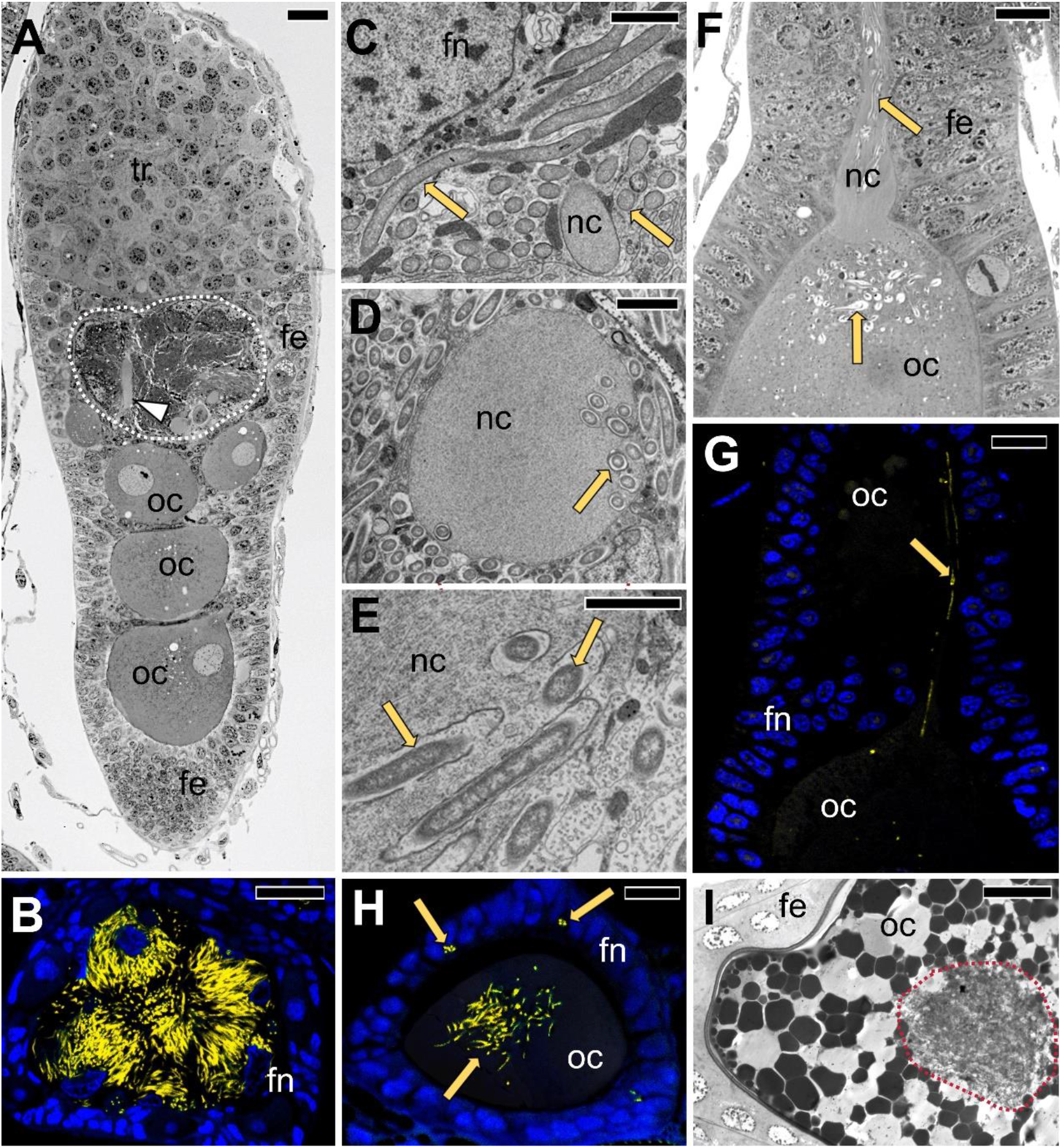
Transovarial transmission of *Sodalis* into the anterior pole of the developing oocyte in representatives of Dictyopharinae subfamily. **A.** Longitudinal section through the ovariole in the previtellogenesis stage. Note numerous *Sodalis* cells in the neck region of the ovariole (area surrounded with white dotted line). *D. pannonica*, LM, scale bar = 20μm. **B.** Cross-section through the neck region of the ovariole filled with prefollicular cells with bacteria *Sodalis* (marked in yellow), *C. krueperi*, confocal microscope, scale bar = 20 μm. **C.** Fragment of the prefollicular cell with *Sodalis* occupying the neck region of the ovariole. *D. multireticulata*, TEM, scale bar = 2 μm. **D.** Cross-section through the nutritive cord surrounded by *Sodalis*. Note *Sodalis* cells in the cytoplasm of the nutritive cord. *C. krueperi*, TEM, scale bar = 2 μm. **E.** The higher magnification of *Sodalis* bacteria migrating via nutritive cord to the oocyte. *C. krueperi.* TEM, scale bar = 2 μm. **F, G.** The transport of *Sodalis* via nutritive cord to the previtellogenic oocyte. *C. krueperi.* **F.**LM, scale bar = 20 μm. **G.** CLM, scale bar = 20 μm. **H.** Accumulation of *Sodalis* cells in the cytoplasm of the anterior region of the previtellogenic oocyte. *C. krueperi*, CLM, scale bar = 20 μm. **I.** Accumulation of *Sodalis* in the cytoplasm of the choriogenic oocyte. *D. multireticulata*, LM, scale bar = 20 μm. Abbreviations: fe – follicular epithelium, fn – the nucleus of a follicular cell, nc or arrowhead – nutritive cord, oc – oocyte, yellow cells/arrows – *Sodalis.*

**Figure 8.**
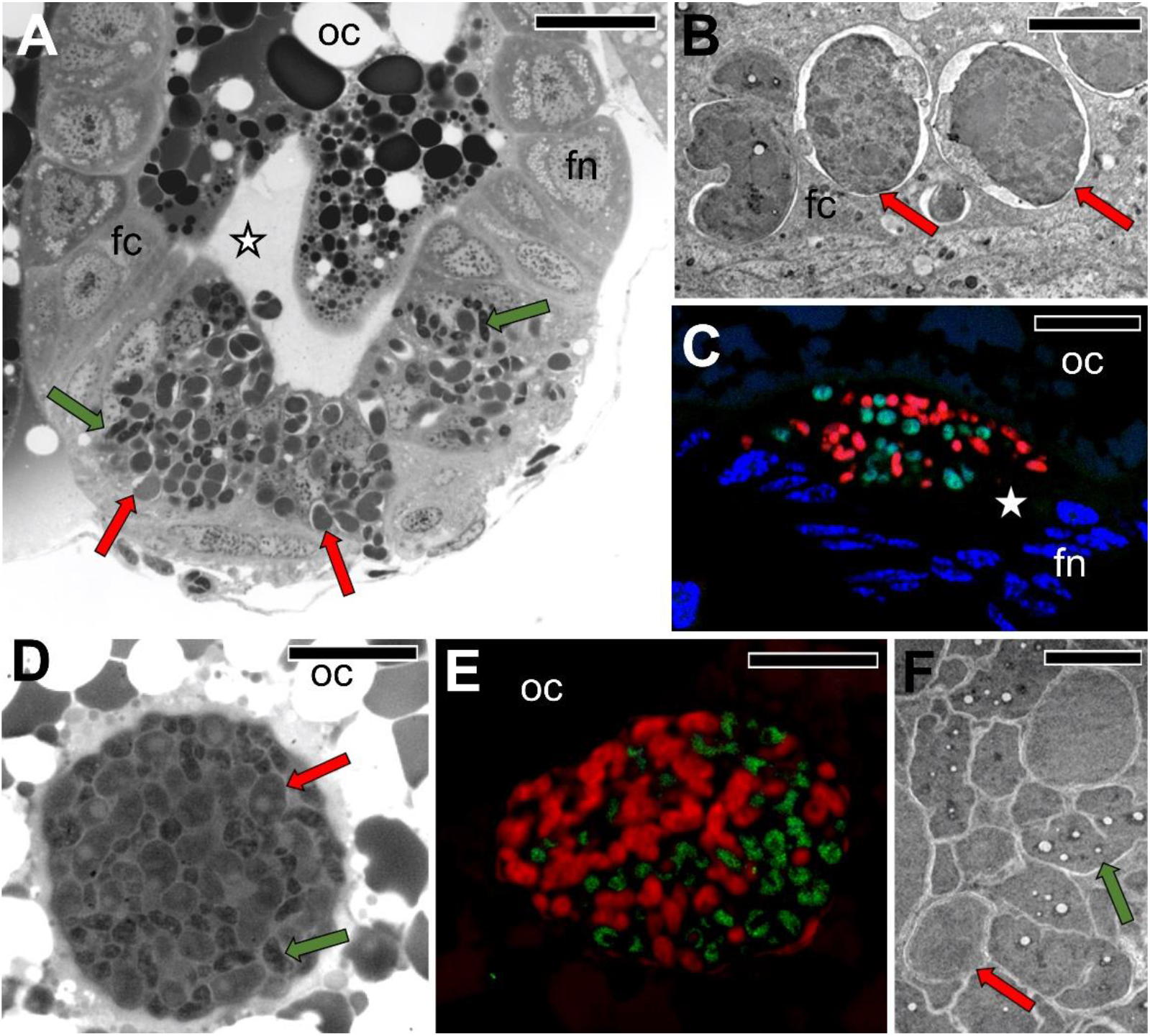
Transovarial transmission of *Sulcia* and *Vidania* into the posterior pole of the terminal oocyte in representatives of Dictyopharinae subfamily. **A.** The migration of *Sulcia* and *Vidania* to the perivitelline space through the follicular epithelium surrounding the posterior pole of the terminal oocyte. *D. multireticulata*, LM, scale bar = 20 μm. **B.** *Vidania* in the cytoplasm of the follicular cell. *D. pannonica*, TEM, scale bar = 2 μm. **C.** Accumulation of *Sulcia* (green) and *Vidania* (red) in the perivitelline space. *C. krueperi.* Confocal microscope, scale bar = 20 μm. **D.** A “symbiont ball” containing *Sulcia* and *Vidania* in the deep depression of the oolemma at the posterior pole of the terminal oocyte. *D. multireticulata*, LM, scale bar = 20 μm. **E.** *In situ* identification of symbionts in the “symbiont ball” in the mature oocyte of *C. krueperi*. CLM, scale bar = 20 μm. **F.** Fragment of “symbiont ball”. *D. multireticulata*. TEM, scale bar = 2 μm. fc – follicular cell, fn – the nucleus of a follicular cell, oc – oocyte, asterisk – perivitelline space, red cells/arrows – *Vidania*, green cells/arrows – *Sulcia*.

In contrast to *Sulcia*, *Sodalis,* and *Arsenophonus,* which do not change their shape substantially during migration, *Vidania* undergoes significant morphological changes. In comparison to lobate *Vidania* cells within bacteriocytes, migrating *Vidania* cells (in the cytoplasm of follicular cells and in the “symbiont ball”) are smaller and more spherical (Figs 6; 8). Transmitting *Vidania* cells resemble those occupying the rectal organ.

## Discussion

### Repeated symbiont replacements shape the symbiosis in Dictyopharidae planthoppers

Amplicon sequencing, metagenomics, and microscopy data agreed that all studied species of Dictyopharidae host heritable nutritional symbionts *Sulcia* and *Vidania*. We know that *Sulcia* infected the common ancestor of all Auchenorrhyncha that lived about 300 MYA, and *Vidania*, together with spittlebug-associated *Zinderia* and leafhopper-associated *Nasuia*, appears to represent a similarly ancient lineage (2, 23, 24). Both of these symbionts are retained in many modern planthoppers (15, 20–22), but so far, their genomes have been characterized in only one other planthopper species, *O. filicicola* OLIH (Cixiidae). We found surprisingly few differences in the symbiont genome organization or contents between Dictyopharidae and OLIH, despite them being separated by about 200 MY of evolution (2). While in Dictyopharidae, symbiont genomes are somewhat smaller than those in OLIH (142-148 kb vs 156 kb for *Sulcia*, 122-126 kb vs 136 kb for *Vidania*), they are co-linear and encode essentially the same set of nutrient biosynthesis genes. *Sulcia*-CALKRU has the most compact genome out of many strains of this symbiont characterized so far. *Vidania* falls among bacteria with the smallest known genomes, alongside other hemipteran symbionts: *Nasuia* (**≥**110 kb), *Tremblaya* (**≥**138 kb), *Carsonella* (**≥**164 kb), and *Hodgkinia* (≥ 144 kb, but note that individual genomes in fragmented complexes can be much smaller) (7, 25–28).

The reduction of symbiont genomes in long-lasting symbiotic interactions is a natural consequence of the lack of DNA repair genes or recombination, allowing for the progressive accumulation of deleterious mutations. Combined with the increase in rates of evolution, this leads to rapid loss of genes and pathways and is thought to negatively affect the function and efficiency of others, including those involved in essential nutrient biosynthesis (29–31). This opens up a path to their complementation or replacement by newly arriving microorganisms, more versatile and efficient. From the evolutionary perspective, both the complementation and replacement of “old” symbionts by “new” ones should allow host insects to refresh and reset their biosynthetic capacity and likely positively influence fitness and perhaps extend ecological niche (7, 23). The radiation of insect clades whose ancestors acquired new symbionts, including cicadas, sharpshooter leafhoppers, aphids, carpenter ants, and others, is suggestive of a competitive edge that the nutritional symbiont acquisition or replacement can provide.

We found that in addition to the two ancestral symbionts, all studied Dictyopharidae species harbor heritable, bacteriocyte-associated enterobacteria, either *Sodalis* or *Arsenophonus,* apparently acquired several times independently. These two microbial clades have repeatedly infected diverse insects including multiple hemipteran lineages. *Sodalis* is known as a nutritional symbiont of blood feeding flies and lice, lygaeid bugs, and weevils (32–34), but a strain has also been isolated from a human wound (35). *Arsenophonus* has assumed different functions in different hosts: some strains are obligatory, heritable nutritional mutualists of blood sucking flies or other insects, others - facultative symbionts, reproductive manipulators, or insect-vectored plant pathogens (2, 36–39). In Auchenorrhyncha, *Sodalis* and *Arsenophonus* usually associate with ancestral symbionts, presumably complementing them (34). For example, they have been reported alongside ancestral symbionts in leafhoppers *Macrosteles laevis* and *Cicadella viridis* and in the spittlebug *Aphrophora quadrinotata* (10, 12, 40). Other times, they appear to have replaced one of the ancient symbionts. For example, Koga and colleagues (12) reported a link between the loss of *Zinderia* and the acquisition of *Sodalis* in modern Philaenini spittlebugs. There is little information on how often such replacements happen; but we think that serial replacements are the most likely explanation for the diversity of *Sodalis*/*Arsenophonus* symbionts within the studied Dictyopharidae. Likewise, Husnik and McCutcheon (26) suggested that this is a plausible explanation for the diversity of enterobacterial symbionts within mealybugs.

Unfortunately, 16S rRNA data provide limited phylogenetic resolution, and the scarcity of genomic references for these broadly distributed and significant symbionts makes it hard to resolve relationships among strains conclusively. However, the distribution of symbiont strains on the host phylogeny, alongside differences in how *Sodalis* in the two subfamilies is transmitted, makes it apparent that the current enterobacterial symbiont distribution across the studied planthopper species is the result of not less than four infection/replacement events.

The diversity of gammaproteobacterial symbionts, including also *Serratia* and *Symbiopectobacterium* in mealybugs, aphids, and other systems, raises questions about the actual dynamic, temporal scales of infections and replacements with these microbes. They can only be addressed using systematic genomics-enabled surveys within insect clades.

### Remarkable conservation of tissue localization and functions in dynamic planthopper symbioses

In all Dictyopharidae species analyzed, as in other planthoppers examined previously, the ancestral symbionts *Sulcia* and *Vidania* are localized in distinct, spatially separated bacteriomes within the insect body cavity (15, 41–43). This contrasts with the situation in Cicadomorpha, where the ancestral symbionts always inhabit the same bacteriome: *Sulcia,* as a rule, occupies bacteriocytes within the outer layer of the bacteriome, whereas the co-residing ancestral symbiont (*Nasuia*, *Zinderia* or *Hodgkinia*) occupies the central portion (11, 27, 44). The differences in the spatial organization of symbiont-housing organs may be due to chance, but could also be related to their mutual relations and nutritional capabilities. In Cicadomorpha, co-primary symbionts are thought to share some cofactors and metabolites, mainly involved in the biosynthesis of energetically expensive amino acids such as methionine or histidine (4), and this could be facilitated by the proximity of bacteriocytes inhabited by different symbionts.

While the ancestral hemipteran symbionts are usually housed in dedicated insect cells – bacteriocytes, the localization of more recently acquired symbionts may differ. Gammaproteobacterial symbionts, including *Arsenophonus* and *Sodalis*, localize differently in different insect hosts. The gammaproteobacterial symbionts of Dictyopharidae, despite their varied origins, are always located within bacteriomes separated from, but adjacent to, those occupied by *Sulcia* and *Vidania*. In other Auchenorrhyncha, they can also occur in separate bacteriomes, or alternatively, colonize distinct bacteriocytes within existing bacteriomes, but they are also observed in the cytoplasm of the same bacteriocytes as ancestral symbionts, or even inside other symbionts’ cells (10, 11, 19). In other insects, they may be dispersed across host tissues other than bacteriocytes, including gut epithelium cells, fat body cells, and hemolymph (45, 46). Interestingly, in some systems, independently acquired microbes tend to inhabit the same tissue compartments, suggestive of pre-adaptations that make it a particularly hospitable place: gammaproteobacterial symbionts of Pseudococcinae mealybugs, always residing inside the cells of the ancient *Tremblaya* symbiont, are a striking example (26).

In females of all Dictyopharidae, similarly to previously studied planthopper species, we observed two morphotypes of *Vidania* occupying distinct bacteriomes in the body cavity and rectal organ. Buchner (15) suggested that symbionts derived from the rectal organ are the infectious form of the same giant lobed symbionts that occur in the bacteriome. Similarly, Bressan and Mulligan (42) hypothesized the functional specialization of *Vidania* bacteriomes into organs containing non-infectious symbiont cells with prominent metabolic functions, and the rectal organs with *Vidania* cells serve for the transmission to the progeny. Future transcriptomic studies should clarify the roles of these morphotypes, but, our microscopic observations agree that *Vidania* cells which are transmitted to the ovary are of rectal organ origin.

In Cicadomorpha, *Sulcia* encodes genes involved in the biosynthesis of seven to eight amino acids, while the companion symbiont is responsible for the provisioning of the remaining two or three (histidine, methionine and sometimes tryptophan). In Fulgoromorpha, the relative contributions of the companion symbionts are reversed: the ability to synthesize arginine, lysine, phenylalanine, and threonine has been lost by *Sulcia* and they are produced by *Vidania* instead. Notably, the near-identity of biosynthetic capabilities in Dictyopharidae and the divergent cixiid, *O. filicicola*, reveals how stable their ancestral nutritional endosymbionts can be over 200 MY of evolution alongside changing gammaproteobacterial partners. The consistent differences in *Sulcia* biosynthetic capabilities between Cicadomorpha and Fulgoromorpha suggest that the two clades have separated soon after the acquisition of the ancestral beta-symbiont – assuming that there was indeed one, as proposed but not unambiguously demonstrated (23).

Later, stochastic or other factors must have caused differential gene and pathway loss among the partner symbionts in the ancestors of the two infraorders.

In planthoppers, at least those studied to date, B-vitamin biosynthesis has been outsourced to additional, gammaproteobacterial symbionts, for example *Purcelliella* in *O. filicicola* or *Arsenophonus* in *Nilaparvata lugens* (47, 48). Indeed, all characterized planthopper-associated *Sodalis* and *Arsenophonus* strains possess largely complete sets of genes involved in riboflavin and biotin production. Additionally, *Sodalis* of *C. krueperi* and *Arsenophonus* of *R. scytha* may be able to synthesize pyridoxine and folate, respectively. A similar biosynthetic potential was found in *Sodalis*-related symbionts of mealybugs (26) and *Arsenophonus* living in symbiotic relations with other insects such as the wasp *Nasonia vitripennis*, and many whiteflies and louse flies (33, 34, 49). Additionally, both *Sodalis* and *Arsenophonus* possess some genes from essential amino acid biosynthesis pathways. Most of them overlap with those retained on the *Vidania* genome. However, in *C. krueperi*, *Sodalis* might complement *Vidania* in the biosynthesis of methionine and in *D. multireticulata, Sodalis* retains two genes necessary in lysine production that underwent pseudogenization in the *Vidania* genome, suggesting that different microbes may complement each other in the nutrient biosynthesis. Such biosynthetic pathway complementarity between the host and symbionts has been demonstrated experimentally in the mealybug system (50) but is likely to be a more universal phenomenon. It would be interesting to explore more broadly whether there is a general trend towards complementarity, how it is affected by serial co-primary symbiont replacements, and whether it can facilitate the replacement of ancient symbionts.

### Newly arriving symbionts follow different strategies of transmission across host generations

The crucial role of nutritional symbionts in host-insect biology is evidenced by their vertical transmission, which ensures the presence of symbionts in subsequent generations. Insects have adopted multiple vertical transmission strategies (16, 17), but the transovarial transmission is probably the most reliable means of providing the full symbiont complement to all offspring. In Auchenorrhyncha, ancient symbionts are transmitted in a consistent, conserved way. In all species studied so far, they cross the follicular cells surrounding the posterior pole of the vitellogenic oocytes and then form the “symbiont ball” near its posterior end (17). Additional symbionts generally follow the same path (11, 44, 51); this includes *Arsenophonus* and *Sodalis* in some of the Dictyopharidae presented here. The discovery that in other Dictyopharidae species, *Sodalis* have adopted a very different transmission strategy, invading the "neck region" of the ovariole from where it is carried to the cytoplasm of previtellogenic oocytes via the nutritive cords, expands our understanding of the diversity of strategies that newly arriving insect symbionts may adopt. The fact that closely related microbes adopted different strategies in related host species is particularly notable. Nonetheless, the spatiotemporal separation of transovarial symbiont transmission is not unique to this insect clade. For example, in the planthopper *Cixius nervosus*, one of its symbionts, unidentified so far, was observed to infect undifferentiated cystocytes, while others full-grown oocytes (17). At the same time, Müller (22), who originally reported different infection strategies in Dictyopharidae, and Buchner (15), who in his pivotal book compared these data against vast amounts of other observations, emphasized their uniqueness.

In other groups of hemipterans and more divergent insects, gammaproteobacterial nutritional symbionts have adopted a wider range of strategies. For instance, in some scale insects, bacteria invade larval ovaries and germ cells at a very early stage, before they differentiate into oocytes and trophocytes, and are later present in all germ cells in the ovariole (17). In some heteropteran bugs but also carpenter ants, symbiotic gammaproteobacteria directly infect early previtellogenic oocytes: infect follicular cells, gather in their cytoplasms, and then enter the oocyte’s cytoplasm via an endocytic pathway. In both cases, initially, symbionts are dispersed in the entire ooplasm, but during vitellogenesis they accumulate and form “symbiont ball” (52, 53). In some host lineages, enterobacterial symbionts have adopted more exotic transmission modes. In *Cicadella viridis* and *Macrosteles laevis*, gammaproteobacterial symbionts (*Sodalis* and *Arsenophonus*, respectively) live inside and are moved to the oocyte within *Sulcia* cells. In the scale insect *Puto superbus*, whole, intact bacteriocytes containing *Sodalis* are transferred into oocytes (10, 40, 54). In turn, in tsetse flies, *Sodalis glossinidius* is vertically carried into developing larva via milk gland secretions (55).

Fitness differences between these alternative modes of transmission are not apparent. Theoretically, the early presence of symbionts in oocytes could contribute to their faster development, but there is no evidence supporting this hypothesis. It is also likely that the transmission modes vary in efficiency, including in the proportion of cells departing from the bacteriomes that arrive within the oocytes, or the energy needed for each symbiont cell to complete the journey. It is also probable that there are differences in the maintenance costs of the mechanisms necessary to move the cells between bacteriomes and oocytes, and later, to the target regions of the developing embryo. Transporting gammaproteobacterial symbionts along the same path as *Sulcia* and *Vidania* could theoretically allow the host to reuse the existing cellular machinery in a relatively efficient manner, but that would depend on the specificity of the mechanisms, something that we have virtually no knowledge of. Yet another aspect relevant to fitness, especially in the long run, is the symbiont bottleneck size, which may affect the strength of genetic drift and the symbiont evolutionary rates (51). However, at the moment, we know little about the magnitude of any such differences, or how important they might be. But ultimately, and most critically, all mechanisms appear to result in reliable transmission of all symbionts to every offspring.

Why do some symbionts adapt different ways of transmission then others, then? We suspect that this is not typically the outcome of selection, but rather a combination of pre-adaptations, existing constraints, and chance events. The transmission strategy, but also final tissue localization, could be related to the microbe’s biology prior to the transition to the endosymbiotic lifestyle. The presence of specific genes or pathways in the microbe’s genome, and mechanisms that direct them towards, or facilitate entry into, certain cell types, are likely to drive the differences (56). At the same time, the re-utilization of existing host-encoded mechanisms, including those for the transmission of older symbionts, is likely to play a significant role, probably increasing over time as symbiont genomes degenerate and erode away. For example, *Spiroplasma* and *Wolbachia* were shown to utilize the receptors involved in the transport of vitellogenin (protein precursors of yolk) into oocytes to enable entry into developing eggs (57, 58). Unfortunately, we can only speculate about the evolutionary processes that have led to this. We suspect that stochastic processes – random changes affecting the affinity between host and symbiont structural molecules at early stages of infection – could play a significant role in determining the way the symbionts end up transmitting.

### How to succeed as a newly arriving heritable symbiont

Transmission between the host generations is a critical challenge for a heritable nutritional endosymbiont, but only one of several facing the microbe transitioning towards obligate symbiosis (59). Yet, despite the multitude of challenges, it appears that successful infections and replacements are relatively common, probably more so than thought until recently (13, 26, 60). Then, what is the recipe for the successful colonization of the host by the new symbiont – one leading to a stable, long-term nutritional endosymbiotic association?

To begin, it seems advantageous for the colonizing microbe to be a pathogen – a product of selection for the ability to gain entry into hosts, overcome or avoid barriers (cuticle) and defenses (behaviour, immune system), and access host resources. Infections would often lead to either disease and death of the host, or alternatively, the clearing of the pathogen by the host immune system. However, low-virulence chronic infections, like those established in some insects by *Sodalis praecaptivus* (61), could provide ample opportunities for gaining entry into oocytes using some existing mechanisms – the first step of transition into a heritable associate. Pathogenic ancestry has been demonstrated for multiple fungal symbionts of cicadas, all hauling from specialized cicada-pathogenic lineages of the genus *Ophiocordyceps* (13), and suspected for bacterial genera that established symbioses repeatedly, including *Sodalis*, *Arsenophonus*, or *Symbiopectobacterium* (37, 60). Although our knowledge of natural strain-level variation is currently limited and this prevents the reconstruction of evolutionary patterns, many symbionts are thought to derive from versatile opportunists similar to *S. praecaptivus*, isolated from a human wound resulting from impalement on a tree branch and thus apparently capable of thriving in very different environments, and falling near the base of the *Sodalis*-allied symbiont phylogeny (35, 62).

Once inside the host, the aspiring nutritional endosymbiont needs to establish residence within specific host organs or tissues and adopt a reliable way of transmitting across host generations. This could be a gradual process with various intermediate stages: for example, the microbe might initially colonize fat body cells that would only later evolve into bacteriocytes (63). As discussed previously, even when related organisms interact, different strategies can be successful. The alternative transmission patterns of *Sodalis*-allied symbiont of Dictyopharidae are not the only example. For instance, in cicadas, following *Hodgkinia* replacement, *Ophiocordyceps* symbionts may reside either in the external layer of the same bacteriomes as the ancient bacterial symbionts, or in a new type of bacteriome (13).

Another aspect important for the long-term success of a recent infection is its effect on the host fitness (64, 65). For the infection to be stable, the symbiont maintenance costs should be counteracted by its positive effects on the host biology. Although stochastic effects such as genetic drift can, in principle, lead to the spread and fixation of a deleterious change (51), the odds of success are clearly greater for the symbiont that positively influences host fitness. The broad metabolic repertoire and efficiency of newly arriving microbes, including their ability to produce essential amino acids and vitamins (13, 34), may constitute obvious benefits to the hosts that lack these nutrients, or are limited by the supply from inefficient, degraded, older symbionts. In some cases, other symbiont-conferred benefits, including the production of defensive compounds (66) may also play a role. Through such effects, the newly arriving symbiont may either complement or replace older symbionts.

Once the symbiont establishes, an unavoidable fate appears to be the degeneration of its genome: rapid pseudogenization and loss of genes and pathways leading to genome shrinkage and overall decrease in function. Some symbionts, for example, *Sulcia* and *Vidania* of dictyopharids, may arrive at the stage where the loss of the remaining small set of critically important genes is essentially halted over tens or hundreds millions of years, but that appears quite uncommon. As they degenerate, symbionts are increasingly dependent on host-encoded mechanisms and products and may ultimately reach a comparable level of dependence as cellular organelles (30, 31). These mechanisms could potentially be reused by different symbionts that co-infect or newly colonize the host. On the other hand, it was shown that two ancient symbionts in a leafhopper rely on different support mechanisms (67). Likewise, the *Sodalis*-allied *Baumannia* symbiont that replaced one of them (*Nasuia*) in a lineage of sharpshooter leafhoppers is supported by a very different set of host genes, suggesting *de novo* evolution and adaptation of the host support mechanisms (18). However, it is plausible that during serial replacements by related bacteria, which seems to be taking place in dictyopharids or mealybugs, it is easier to repurpose the existing symbiont support mechanisms. Once again, we suspect that both the timing and order of gene loss and the recruitment of specific host support mechanisms may be a largely stochastic process, with chance events such as random mutations and genetic drift determining the outcomes. The microbe’s versatility in utilizing host mechanisms may increase its odds of establishing and surviving for a long time as a key part of a successful two- or multi-partite symbiosis. But regardless, once the symbiont establishes within the host, it seems unable to escape the path of genomic degradation and increasing dependence which (23) graphically described as “the rabbit hole”.

The list of microbes that are known to have survived as endosymbionts for an evolutionarily significant amount of time is not very long, and does include the ancestral symbionts of various sap-feeder and blood-feeding insect clades. A large proportion of them, including *Buchnera*, *Baumannia*, *Blochmannia*, and *Wigglesworthia*, have been recently identified as members of the *Sodalis* clade (62). We do not understand what has allowed these symbionts to reach the stable stage (68) rather than degenerate away. But between the high mutational pressure, increasing with progressing genome degeneration, and the constant exposure to new pathogenic and opportunistic microbes, it is becoming increasingly apparent that the majority of gammaproteobacterial symbioses are relatively short-lived. Except for the selected few, the vast majority of such symbionts will succumb to the cycle of serial symbiont replacements.

## Material and Methods

### Study species

We investigated symbioses in seven planthopper species from two subfamilies within the family Dictyopharidae. From subfamily Dictyopharine, we characterized four species: *Callodictya krueperi* (Fieber, 1872), *Dictyophara pannonica* (Germar, 1830), *Dictyophara multireticulata* Mulsant & Rey, 1855, *Dictyophara europaea* (Linnaeus, 1767). From Orgeriinae, we studied three species: *Ranissus (Schizorgerius) scytha* (Oshanin, 1913), *Ranissus (Ranissus) edirneus* (Dlabola, 1957) and *Parorgerius platypus* (Fieber, 1866). Adult specimens were sampled from a single population of each species in either Bulgaria or Poland between 2016 and 2018 (Dataset Table S1), preserved in 80% ethanol or glutaraldehyde, and stored at 4 °C until processing. Representative specimens from each species were identified based on morphological characteristics, and identities confirmed using marker gene sequencing.

### Amplicon-based microbiome screen

#### Library preparation and sequencing

To obtain the preliminary picture of the microbiomes in the seven experimental species, we sequenced amplicons for the V4 hypervariable region of the bacterial 16S rRNA gene, simultaneously confirming the specimen identity by sequencing amplicons for partial COI genes. DNA extracted using Bio-Trace Extraction Kit (Eurx, Gdańsk, Poland) from dissected insect abdomens for up to three individual insects per species (plus negative controls), was used for amplicon library preps following a modified two-step PCR library preparation protocol provided by (69). In the first round of PCR, we amplified two marker regions of interest, using template-specific primers 515F/806R (70, 71) and COIBF3//COIBR2 (72) with Illumina adapter tails. The PCR products were purified using SPRI magnetic beads and then used as the template for the second, indexing PCR reaction. Pooled libraries were sequenced on an Illumina MiSeq v3 lane (2 × 300 bp reads) at the Institute of Environmental Sciences of Jagiellonian University. The primer sequences and detailed protocols for amplicon library preparation are provided in Dataset Table S2.

#### Analyses of amplicon sequencing data

We processed 16S rRNA and COI genes amplicon data using mothur v. 1.43.0 (73) following a pipeline detailed in the Supplementary Material. Initially, all amplicon datasets were split into bins corresponding to the two target genes based on primer sequences, and each bin was analyzed separately. For both bins, we assembled forward and reverse reads into contigs, which were then quality-filtered. The contigs were then de-replicated, and the sequences that occurred only once in the dataset (singletons) were discarded. Then, we aligned contigs against the corresponding reference database, removing those that did not align properly. For analysis of 16S rRNA gene sequences, the aligned sequences were screened for chimeric sequences using UCHIME, and then classified by taxonomy. Finally, the sequences were clustered at 97% identity level using the nearest-neighbor algorithm and divided into OTUs.

### Metagenomic library preparation and sequencing

We sequenced bacteriome metagenomic libraries for three species: *C. krueperi* (specimen ID: CALKRU), *D. multireticulata* (DICMUL), and *R. scytha* (RANSCY). We extracted DNA from dissected bacteriomes of individual insects using a Sherlock AX DNA extraction kit (A&A Biotechnology, Gdynia, Poland). After fragmentation using Covaris E220 sonificator, we used it for metagenomic library preparation using NEBNext Ultra II DNA Library Prep Kit for Illumina (BioLabs, New England), with the target insert length of 350 bp. The library pool, including three target species and other samples, was sequenced on Illumina HiSeq X SBS lane by NGXBio (San Francisco, CA, U.S.A.).

### Metagenome characterization and symbiont genome annotation

Metagenomic reads were filtered by quality scores using ‘iu-filter-quality-minoche’ program included in illumina-utils software v1.4.4 (74) with default parameters, as described by (75). Contigs were assembled using Megahit v1.2.9 (k-mer 255, min contig size 1000) (76). Because of the known issue of index swapping that occurs during cluster formation and sequencing on Illumina platforms (77, 78) which can lead to cross-contamination among samples in multiplexed lanes, we filtered the resulting assemblies for cross-contamination. We discarded all contigs that had >10X greater coverage estimated based on strictly mapped reads from another library with overlapping indices than based on reads from the same library, after confirming that most of the removed low-coverage contigs indeed perfectly matched high-coverage microbial genomes in these libraries.

We identified symbiont contigs using Blastn and tBlastx searches against a custom database containing genomes of multiple known insect symbionts, verifying the identifications using coverage and GC contents information (computed using BBTools v. 38.78). Then, for *Sulcia* and *Vidania* contigs, we confirmed their circularity and contiguity by read mapping and visualization on Tablet v. 1.20.12.24 (79) and the presence of overlapping ends. We re-arranged the circularized genomes to ensure the same orientation and start position as in those published previously. *Arsenophonus*, *Sodalis* and *Wolbachia* genomes were represented by multiple contigs, and we did not attempt to close them.

The genomic contigs of the three latter symbionts were annotated using prokka v.1.14.6 (80), with the default parameters. For *Sulcia* and *Vidania*, prokka but also Interproscan and GhostKOALA annotation attempts left multiple obvious gaps and unannotated or hypothetical proteins, and hence we decided to annotate them using a custom Python script, modified from (27). Annotation was conducted by recursive searches for a manually curated set of alignments of protein-coding, rRNA and ncRNA genes from previously characterized *Vidania* or *Sulcia* lineages using HMMER 3.1b2 (81). Any open reading frames of at least 300 nucleotides that had not been annotated by the script were manually searched using hmmer and blastx/tblastx against UniProt and NCBI databases. All genes annotated as “hypothetical” or unannotated were carefully manually compared against the top hits in other microorganisms using blastp (https://blast.ncbi.nlm.nih.gov) and HMMER 3.3 (https://www.ebi.ac.uk/Tools/hmmer), resulting in the discovery of additional genes. Reference-based annotations of rRNA genes were supplemented by rRNA searches using RNAmmer v. 1.2 (82), and tRNA searches using tRNAscan-SE v. 1.4 (83). For all genes, we aligned all copies classified as functional using mafft v. 7.221 (84). In case of protein-coding genes, alignments were conducted in protein space and reverse-translated to nucleotide space.

The taxon-annotated GC-coverage plots for symbiont contigs were drawn using R v. 4.0.2 (R Development Core Team) with ggplot2 package (85). Genomes were visualized using DNAPlotter GUI. In order to reconstruct amino acid and B vitamin biosynthetic pathways, we translated circular *Sulcia* and *Vidania* genomes as well as *Sodalis*, *Arsenophonus* and *Wolbachia* contigs to amino acids and annotated them against KEGG with GhostKOALA (genus_prokaryotes) (86). After that, the presence of genes involved in biosynthetic pathways was checked manually.

### Cloning, amplification, and phylogenetic analyses

For the four species for which metagenomes were not sequenced (*D. pannonica*, *D. europaea*, *R. edirneus* and *P. platypus*), we obtained full-length 16S rRNA gene sequences of symbionts through molecular cloning in *Escherichia coli* cells as described previously (43). We also PCR-amplified planthopper mitochondrial *CytB* and nuclear 18S and 28S rRNA genes (details in Dataset Table S2). Purified PCR products were Sanger-sequenced by Genomed S.A. (Warsaw, Poland). Trimmed reads were merged into contigs, aligned, and alignments inspected using CodonCode Aligner v. 8.0.2 (CodonCode Corp., Centerville, USA).

We conducted phylogenetic analyses of concatenated insect marker genes and bacterial 16S rRNA genes, including those obtained from metagenomes using MEGA 7 software (87) using Maximum Likelihood algorithm assuming the GTR (for insect genes) and GTR+GAMMA (for bacterial genes) models and with 1000 bootstrap replicates.

### Microscopy

#### Light (LM) and electron (TEM) microscopy

Partially dissected abdomens of females of each of the seven species were fixed in 2.5% glutaraldehyde solution in 0.1 M phosphate buffer (pH 7.4) in the field. In the laboratory, after washing with the same buffer with the addition of sucrose (5.8%), they were postfixed in 1% solution of osmium tetroxide, dehydrated in ethanol and acetone series, and embedded in epoxy resin Epon 812 (SERVA, Heidelberg, Germany). For histological analyses, the resin blocks were cut into serial, semithin sections (1 μm thick), stained in 1% methylene blue in 1% borax, and observed under the Nikon Eclipse 80i light microscope. Ultrathin sections (90 nm thick) for ultrastructural analyses were contrasted with lead citrate and uranyl acetate and observed under the JEOL JEM 2100 electron transmission microscope.

#### Fluorescence microscopy

For fluorescence *in situ* hybridization (FISH), insects preserved in 80% ethanol were rehydrated and postfixed in 4% paraformaldehyde for 2 hours. Then, the material was dehydrated again in the increasing concentration of ethanol and acetone and embedded in the Technovit 8100 resin (Kulzer, Wehrheim, Germany). Resin blocks were cut into semithin sections (1 μm thick) and hybridized overnight at room temperature with symbiont-specific probes. The probe details and hybridization reaction conditions are provided in Suppl. Table S2. After hybridization, the slides were washed in PBS, dried, covered with ProLong Gold Antifade Reagent (Life Technologies), and examined using a confocal laser scanning microscope Zeiss Axio Observer LSM 710.

## Author Contributions

AM and PŁ designed research, AS conducted the sampling, AM, DCF, MK, TS and PŁ performed research and analyzed data, AM, DCF and PŁ wrote the paper.

## Acknowledgments

We thank John McCutcheon for permission to use the facilities at the University of Montana, advice, and valuable comments, Ada Jankowska, Monika Prus, and Mateusz Buczek for laboratory assistance, Marcin Walczak for providing *D. europaea* specimens, Gernot Kunz and Ilia Gjonov for insect images. This project was supported by the Polish National Science Centre grants 2017/26/D/NZ8/00799 (to A.M.) and 2018/30/E/NZ8/00880 (to P.Ł.) as well as Polish National Agency for Academic Exchange grant PPN/PPO/2018/1/00015 (P.Ł.). The open-access publication of this article was funded by the Priority Research Area BioSunder the program “Excellence Initiative – Research University” at the Jagiellonian University in Krakow

## Supplementary information and data availability

Supplementary Information file includes nine Supplementary Figures and the details of amplicon analysis workflow. Supplementary Tables file includes twelve Supplementary Tables. Accession numbers for all sequencing datasets are listed in Supplementary Table S11.

