## Supporting Information for "Alternative transmission patterns in independently acquired nutritional co-symbionts of Dictyopharidae planthoppers"

### I. Supplementary figures

Figure S1. The relative abundance of symbionts in the examined Dictyopharidae species
Figure S2. ML phylogeny of *Sulcia* based on 16S rRNA gene sequences
Figure S3. ML phylogeny of *Vidania* based on 16S rRNA gene sequences
Figure S4. ML phylogeny of *Sodalis* based on 16S rRNA gene sequences
Figure S5. ML phylogeny of *Arsenophonus* based on 16S rRNA gene sequences
Figure S6. The alignment of planthopper *Sulcia* genomes
Figure S7. The alignment of planthopper *Vidania* genomes
Figure S8. The summary of nutritional symbiont contribution to amino acid and B-vitamin biosynthesis in different Auchenorrhyncha
Figure S9. *Wolbachia* symbiont cells within bacteriocytes of Dictyopharidae species (TEM)

### II. Bioinformatic pipelines

The analysis of COI amplicon data
The analysis of 16S rRNA amplicon data

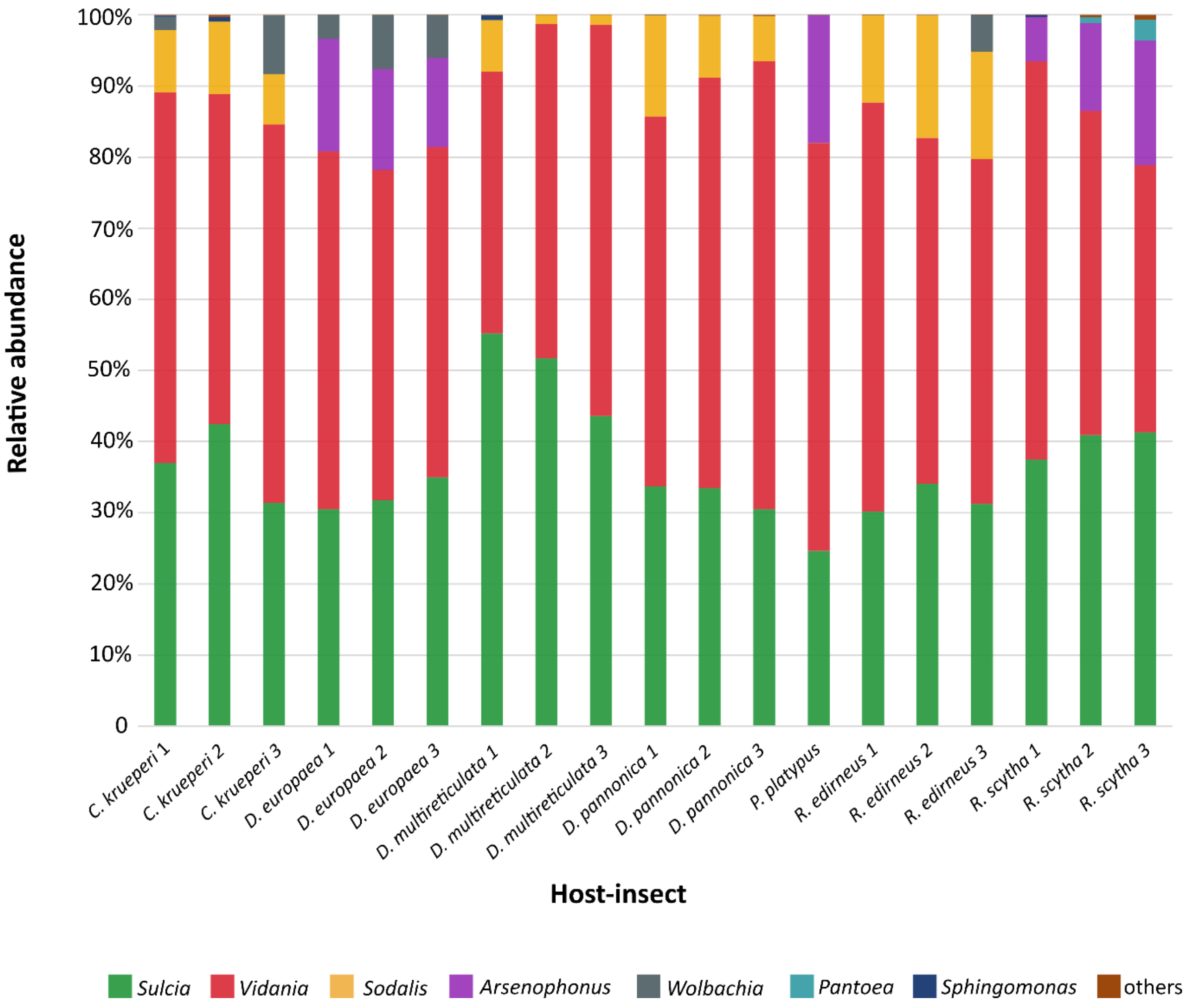


**Figure S1.** The relative abundance of the dominant taxa of symbiotic bacteria in 19 experimental specimens of seven Dictyopharidae species.

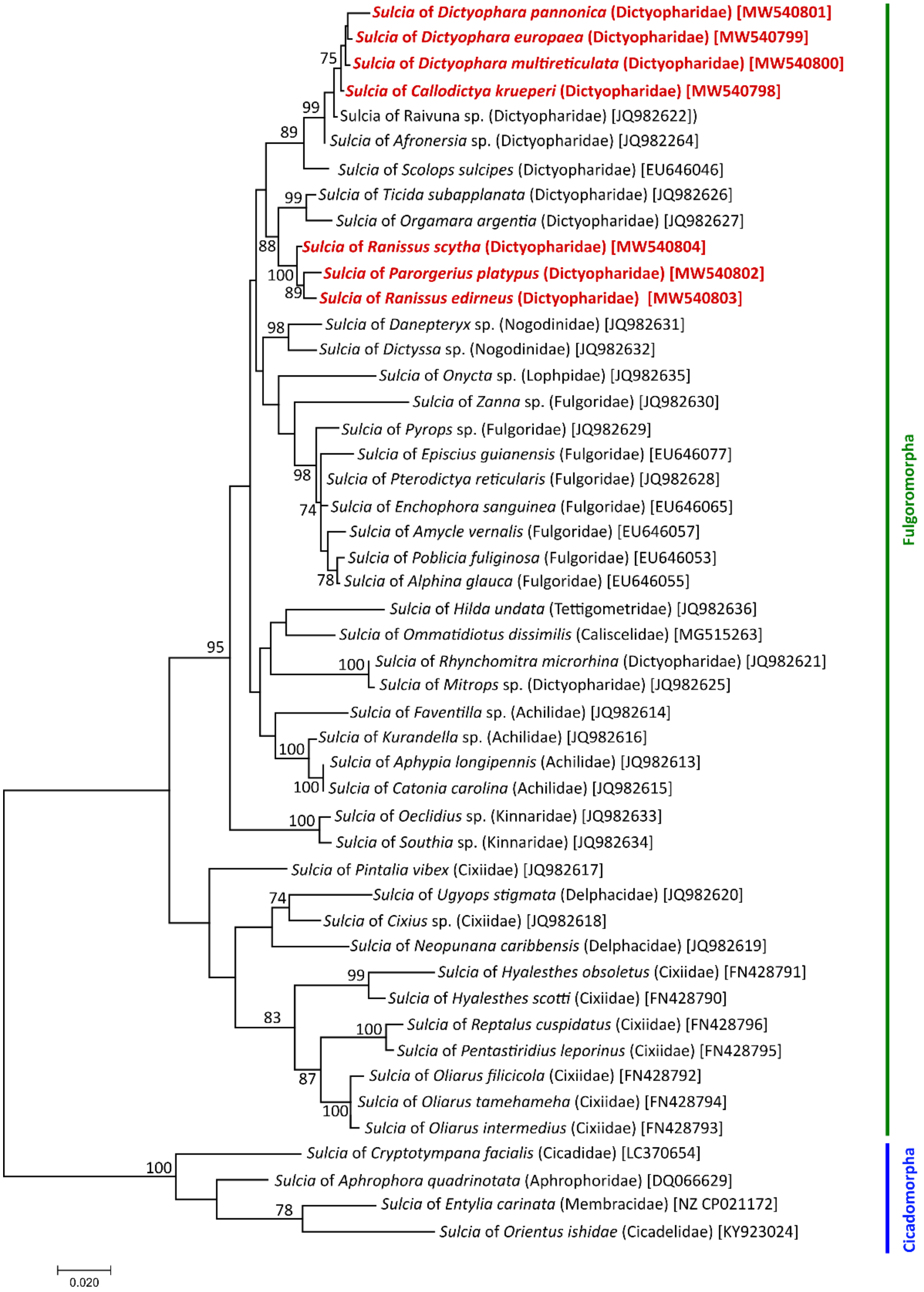


**Figure S2.** ML phylogeny of *Sulcia* strains from diverse planthoppers, based on full-length sequences (1373 shared nucleotide positions) of 16S rRNA gene. Only bootstrap values above 70% are shown. Sequences from the seven Dictyopharidae species studied here are highlighted using the colored font.

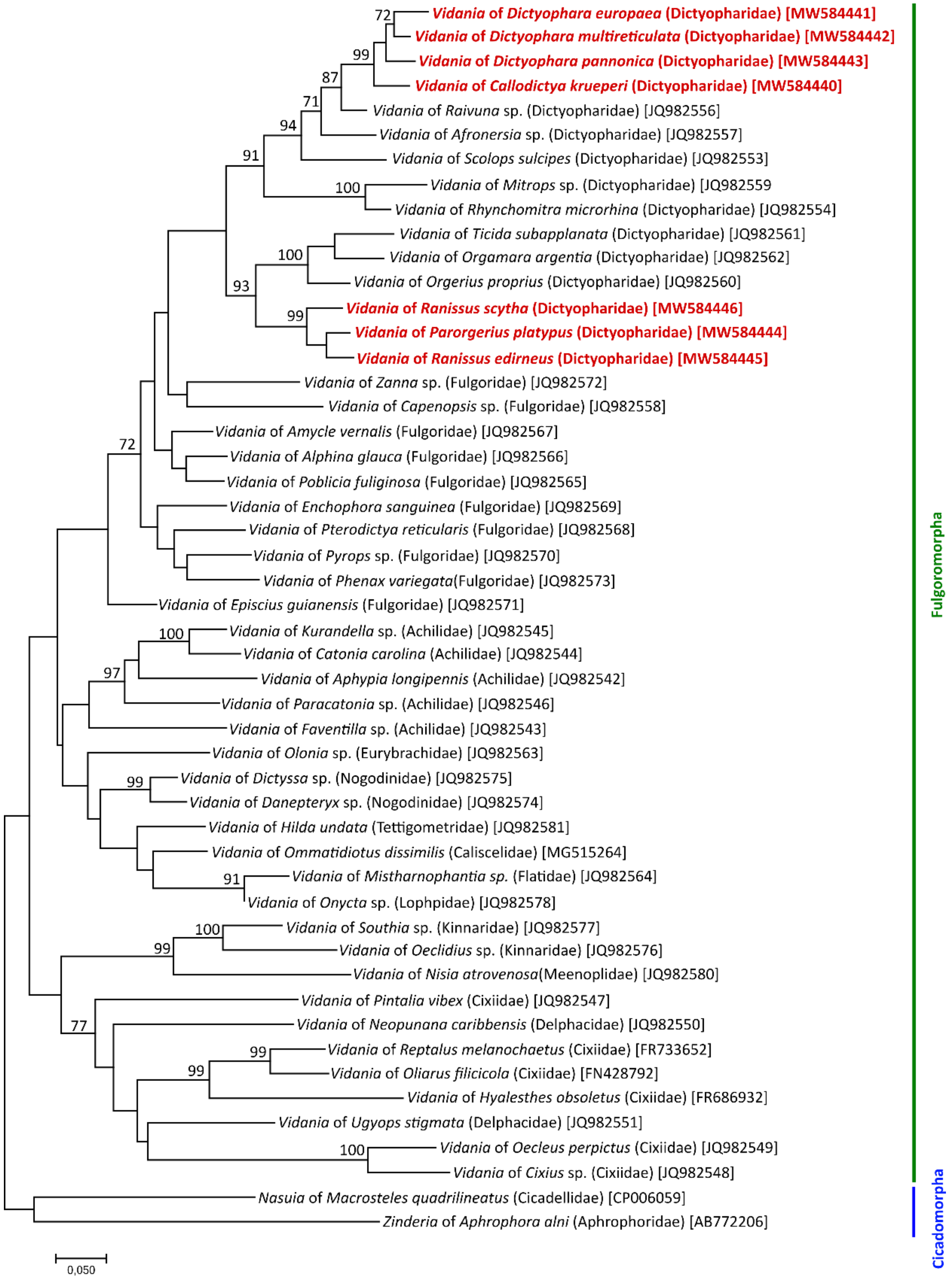


**Figure S3.** ML phylogeny of *Vidania* strains from diverse planthoppers based on full-length sequences (1530 shared nucleotide positions) of 16S rRNA gene. Only bootstrap values above 70% are shown. Sequences from the seven Dictyopharidae species studied here are highlighted using the colored font.

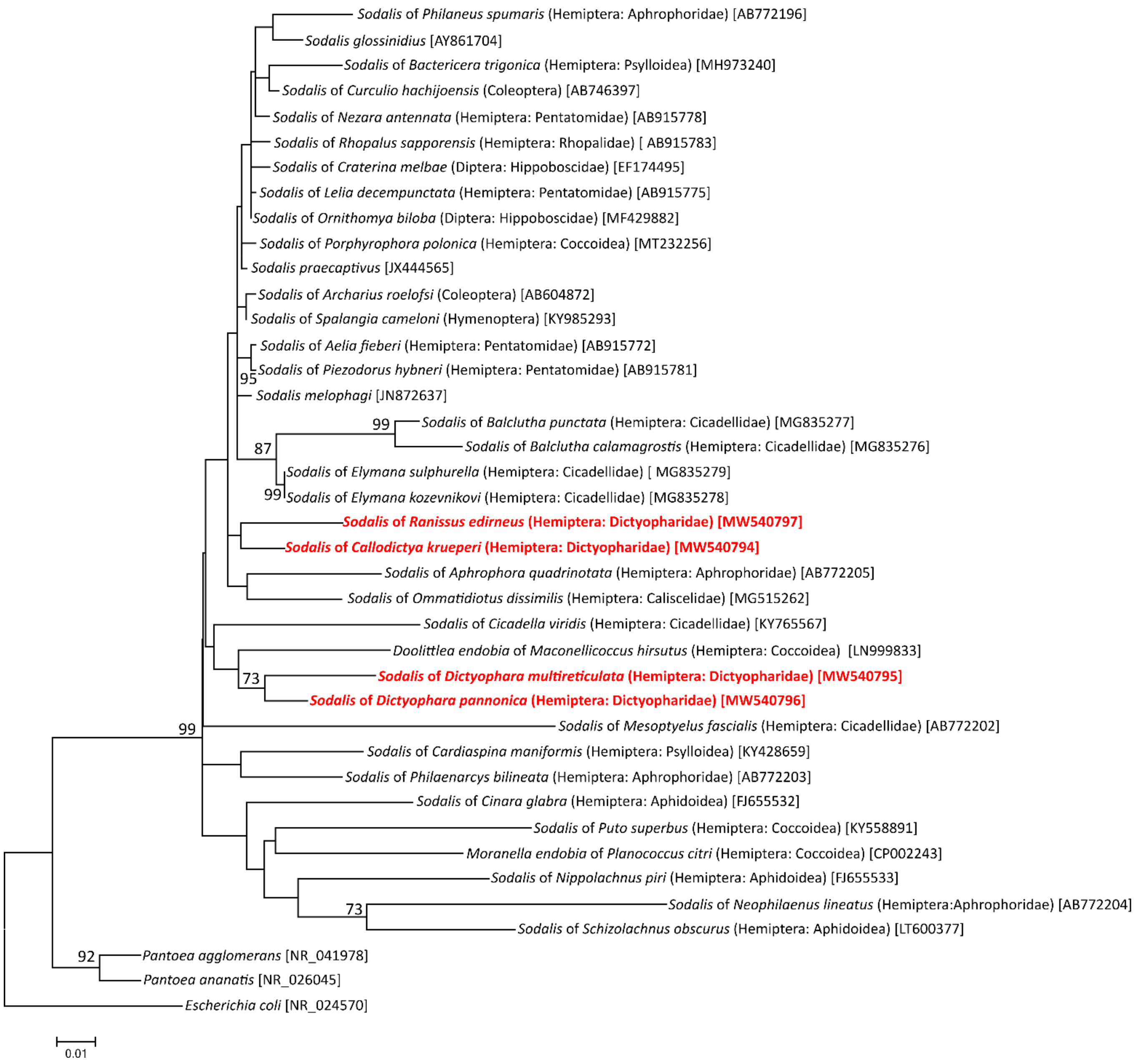


**Figure S4.** ML phylogeny of *Sodalis* strains from different hosts based on 16S rRNA gene sequences. Only bootstrap values above 70% are shown.

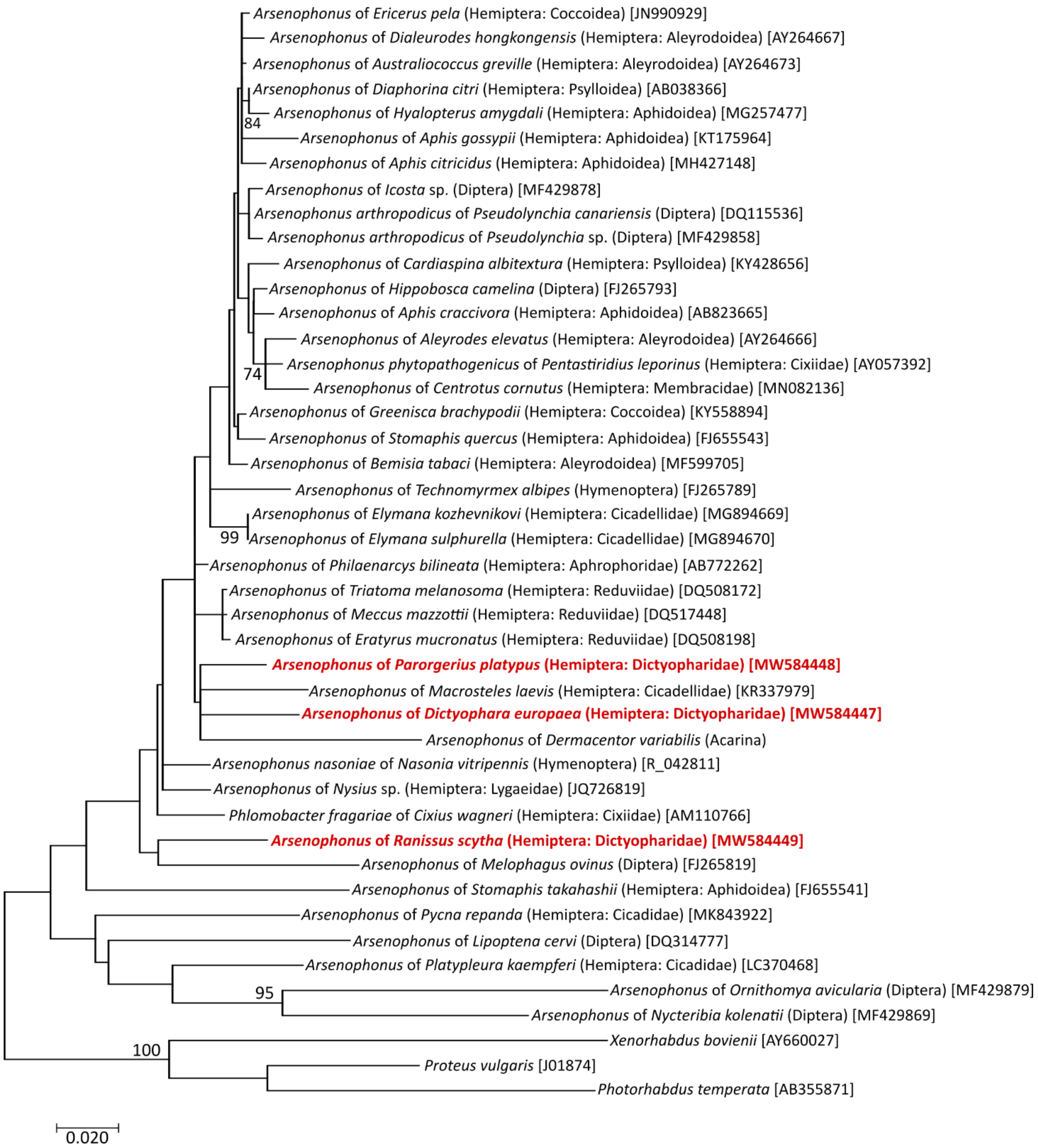


**Figure S5.** ML phylogeny of *Arsenophonus* strains from different hosts based on 16S rRNA gene sequences. Only bootstrap values above 70% are shown.

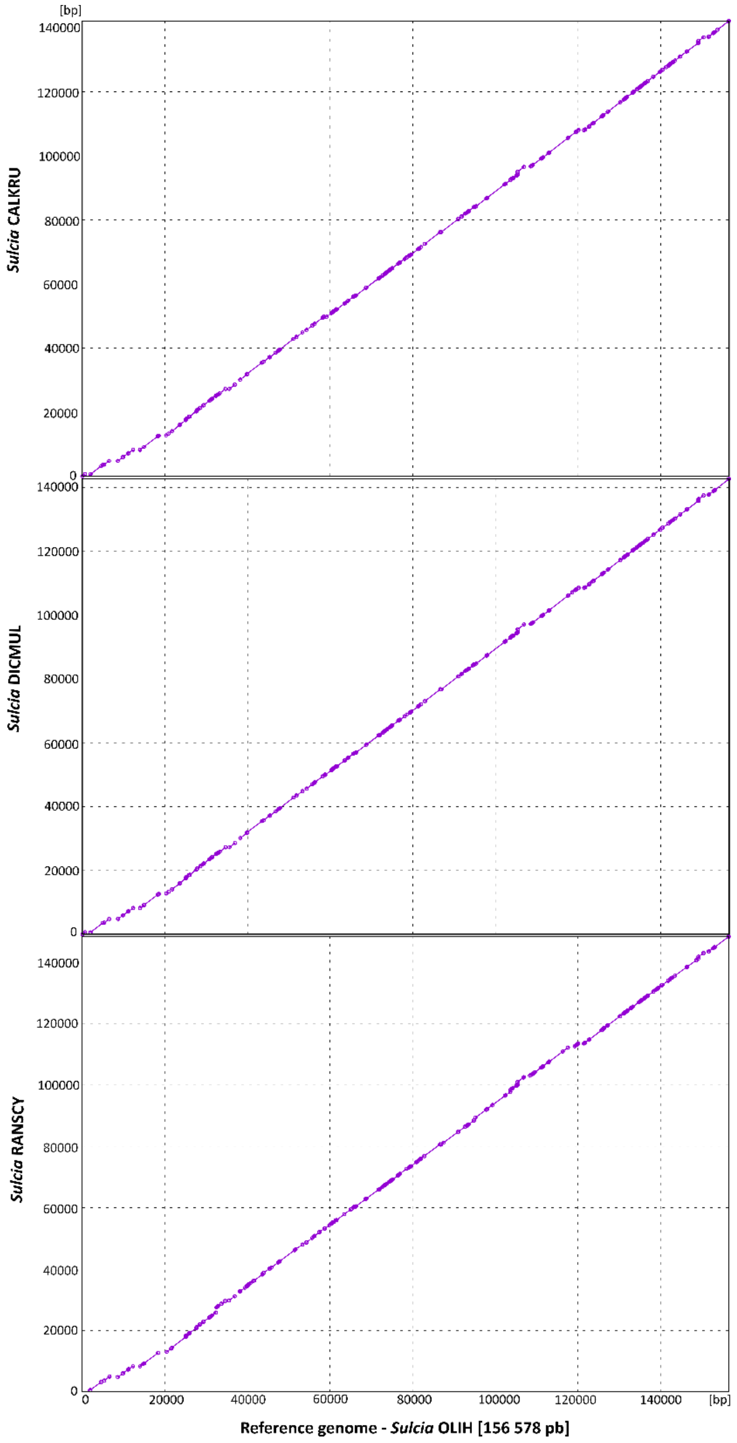


**Figure S6.** PROMER alignments of final *Sulcia* genomes from three Dictyopharidae species: *Callodictya krueperi* (CALKRU), *Dictyophara multireticulata* (DICMUL), and *Ranissus scytha* (RANSCY) against the *Sulcia* genome from *Oliarus filicicola* (OLIH; family Cixiidae).

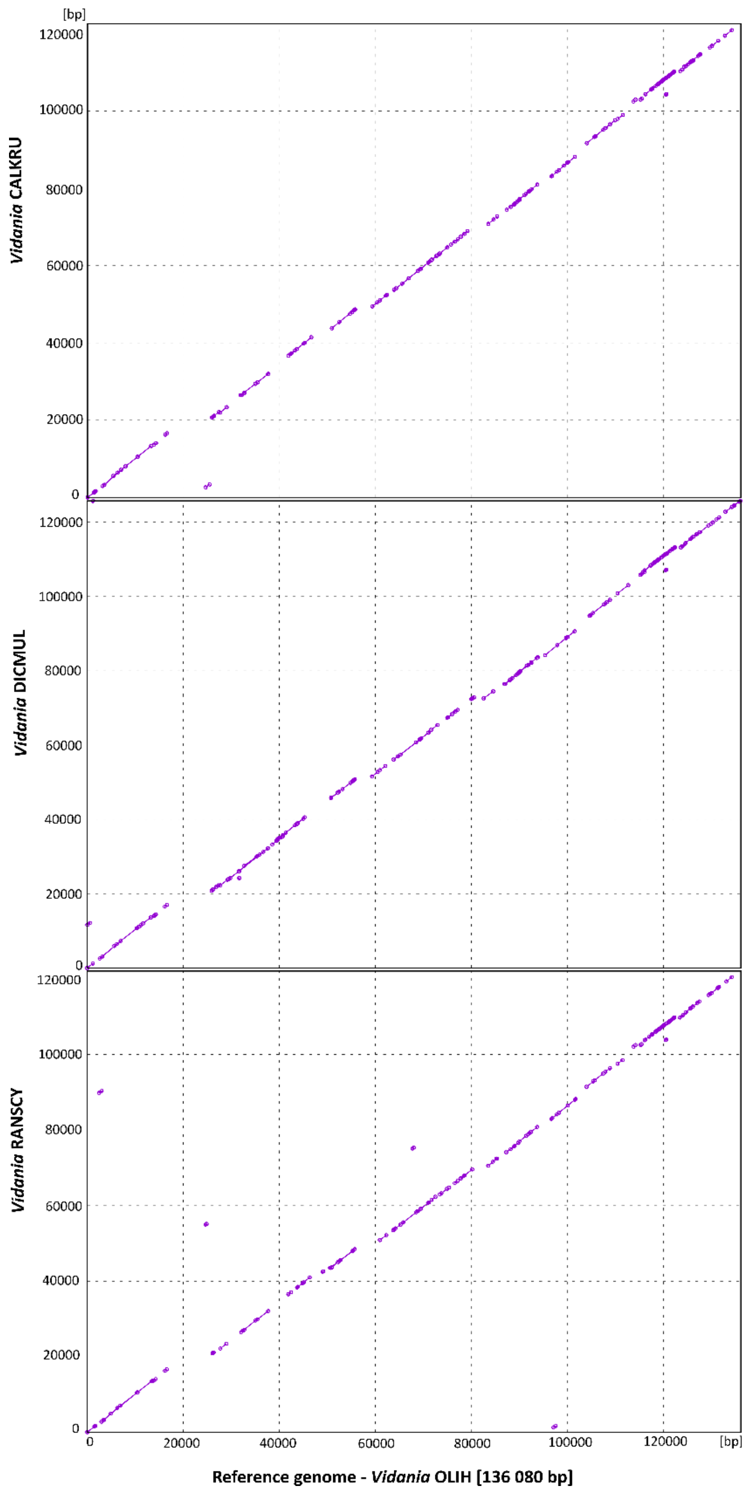


**Figure S7.** PROMER alignments of final *Vidania* genomes from three Dictyopharidae species: *Callodictya krueperi* (CALKRU), *Dictyophara multireticulata* (DICMUL), and *Ranissus scytha* (RANSCY) against the *Vidania* genome from *Oliarus filicicola* (OLIH; Cixiidae).

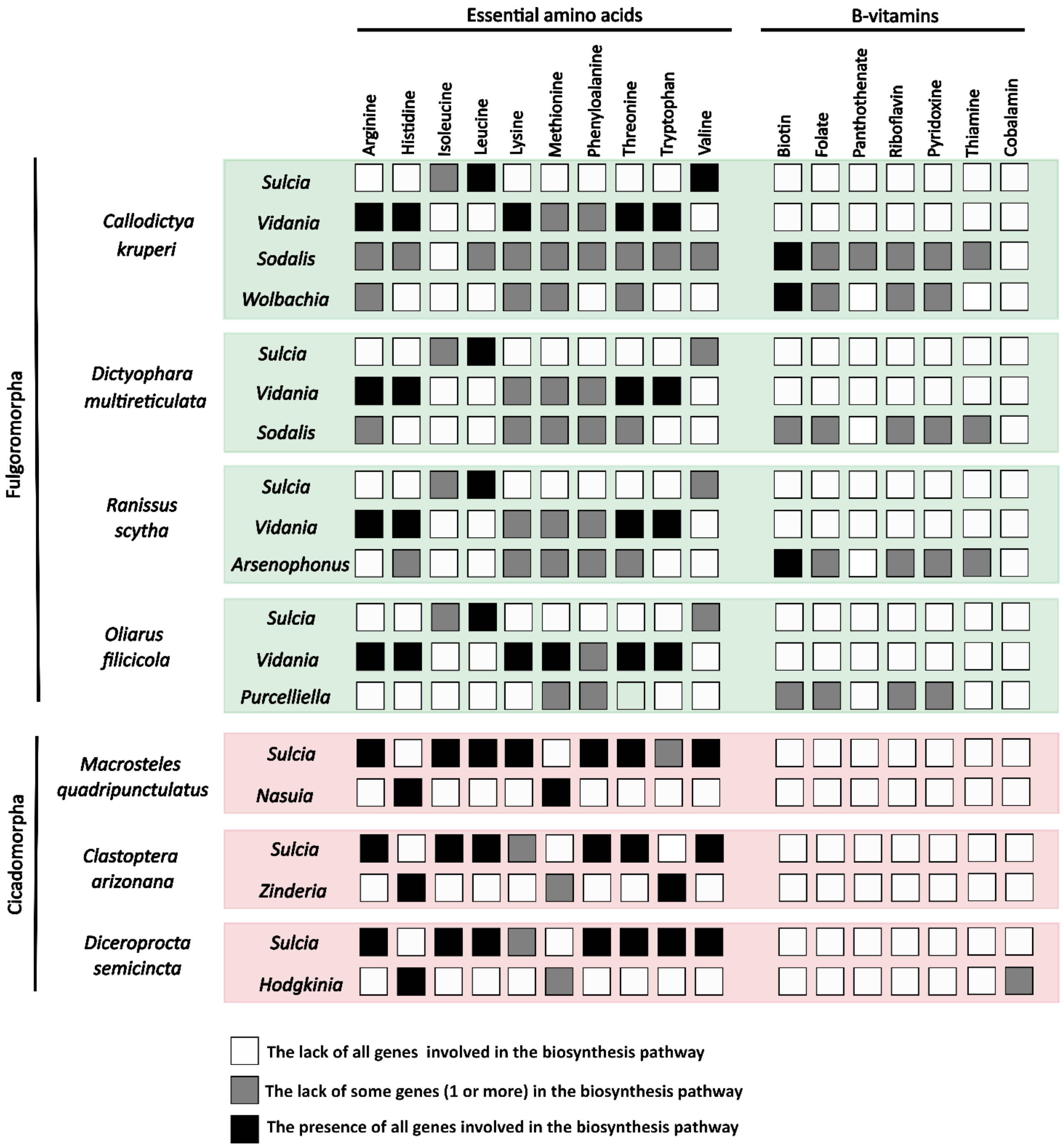


**Figure S8.** The overview of the contributions of endosymbionts in different Auchenorrhyncha to the amino acid and B-vitamin biosynthesis.

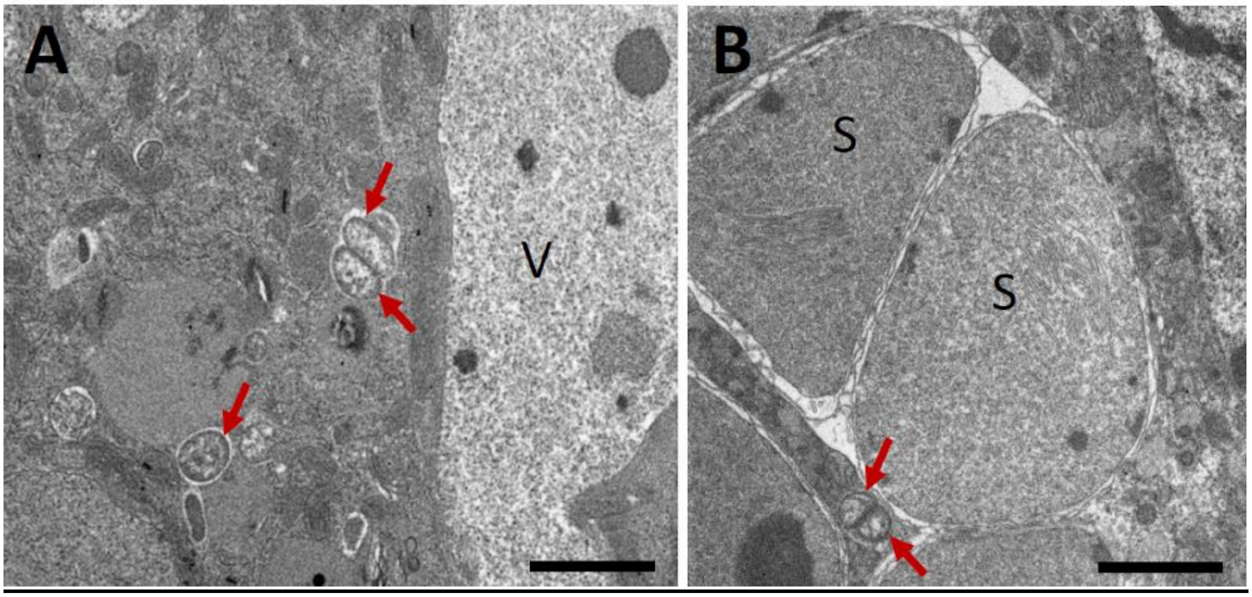


**Figure S9. *Wolbachia* symbiont cells within the bacteriocytes in two Dictyopharidae species.** **A.** *Wolbachia* cells (red arrows) in the cytoplasm of the *Vidania* (V) bacteriocyte from *Callodictya krueperi* **B.** *Wolbachia* cells (red arrows) in the cytoplasm of the *Sulcia* (S) bacteriocyte from *Dictyophara europaea*. TEM, scale bar = 2 µm.

**Bioinformatic pipeline for the analysis of COI amplicon data**

COI amplicon data analysis was conducted using mothur v. 1.43.0. The following commands were used.

```
######## Setting working directories
set.dir(input=/home/anna.michalik/Fulgoromorpha/COI/COI2,
output=/home/anna.michalik/Fulgoromorpha/COI/COI2)

######## Creating the file with a list of libraries and files
make.file(inputdir=/home/anna.michalik/Fulgoromorpha/COI/COI2, type=fastq,
prefix=DIC_COI)

######## Assembling forward and reverse reads into contigs
make.contigs(file=DIC_COI.files)

######## Quality-trimming sequences
trim.seqs(fasta=DIC_COI.trim.contigs.fasta, oligos=primers_to_trim.oligos,
minlength=250, maxlength=500, maxambig=0, maxhomop=10, pdiffs=2)

######## Listing sequences in trimmed fasta file, extracting corresponding
sequence IDs from the group file
list.seqs(fasta=DIC_COI.trim.contigs.trim.fasta)
get.seqs(accnos=DIC_COI.trim.contigs.trim.accnos, group=current)

######## Getting basic information about the sequences (e.g., median length,
the number of ambiguous bases)
summary.seqs(fasta=current)

######## Checking the number of reads in libraries before and after quality
trimming
count.groups(group=DIC_COI.contigs.groups)
count.groups(group=DIC_COI.trim.contigs.fasta)

######## Picking unique sequences
unique.seqs(fasta=DIC_COI.trim.contigs.trim.fasta)

######## Generating a unique sequence table, with info on the abundance of each
unique sequence in each library
count.seqs(name=DIC_COI.trim.contigs.trim.names,
group=DIC_COI.contigs.pick.groups, compress=f)

######## Discarding singleton sequences
split.abund(fasta=current, count=current, cutoff=1)

######## Getting basic information about the sequences
summary.seqs(fasta=DIC_COI.trim.contigs.trim.unique.abund.fasta)
```

```
######## Aligning sequences against a custom database containing reference COI
sequences of insects
align.seqs(fasta=DIC_COI.trim.contigs.trim.unique.abund.fasta,reference=COIrefe
rence2.fas)

######## Getting the basic information about alignment
summary.seqs(fasta=current)

######## Removing unaligned or poorly aligned sequences
screen.seqs(fasta=DIC_COI.trim.contigs.trim.unique.abund.align,
count=DIC_COI.trim.contigs.trim.abund.count_table, minlength=400)

######## Removing any columns containing only gap characters
filter.seqs(fasta=DIC_COI.trim.contigs.trim.unique.abund.good.align,vertical=T,
trump=.)

######## Changing file names
rename.file(fasta=current, count=current, prefix=COI)

######## Computing pairwise distance matrix
dist.seqs(fasta=DIC_COI.fasta, cutoff=0.1)

######## Clustering OTUs
cluster(column=current, count=DIC_COI.count_table, cutoff=0.03)

######## Assigning sequences to OTUs based on 97% identity
bin.seqs(list=current, fasta=current, label=0.03)

######## Creating a 97% OTU table
make.shared(list=current, count=current, label=0.03)
```

### Analysis of 16S rRNA amplicon data

The analysis of amplicon data for the V4 region of the 16S rRNA gene was conducted using mothur v. 1.43.0. The following commands were used.

```
######## Setting working directories
set.dir(input=/home/anna.michalik/Fulgoromorpha/DIC2/16Samplicon_analysis,
output=/home/anna.michalik/Fulgoromorpha/DIC2/16Samplicon_analysis)

######## Creating the file with a list of libraries and files
make.file(inputdir=/home/anna.michalik/Fulgoromorpha/DIC2/16Samplicon_analysis,
type=fastq, prefix=16S)

######## Assembling forward and reverse reads into contigs
```

```
make.contigs(file=16S.files)

######## Quality-trimming of sequence data
trim.seqs(fasta=16S.trim.contigs.fasta, oligos=primers_to_trim.oligos,
minlength=250, maxlength=500, maxambig=0, maxhomop=10, pdiffs=2)

######## Listing sequences in trimmed fasta file, extracting corresponding
sequence IDs from the group file
list.seqs(fasta=16S.trim.contigs.trim.fasta)
get.seqs(accnos=16S.trim.contigs.trim.accnos, group=current)

######## Getting the basic information about the sequences
summary.seqs(fasta=current)

######## Checking the numbers of reads in libraries before and after quality
trimming
count.groups(group=16S.contigs.groups)
count.groups(group=16S.trim.contigs.fasta)

######## Identifying unique sequences
unique.seqs(fasta=16S.trim.contigs.trim.fasta)

######## Generating a unique sequence table, with info on the abundance of each
unique sequence in each library
count.seqs(name=16S.trim.contigs.trim.names, group=16S.contigs.pick.groups,
compress=f)

######## Discarding singleton sequences
split.abund(fasta=current, count=current, cutoff=1)

######## Getting basic information about sequences
summary.seqs(fasta=16S.trim.contigs.trim.unique.abund.fasta)

######## Aligning sequences against a SILVA database
align.seqs(fasta=16S.trim.contigs.trim.unique.abund.fasta,
reference=/mnt/matrix/symbio/db/silva.nr_v132.align, processors=20)

# * align.seqs(fasta=16S.trim.contigs.trim.unique.abund.fasta,
reference=/home/anna.michalik/Fulgoromorpha/DIC2/16Samplicon_analysis/new_inter
nal_reference.fas, processors=20)
# * Note: We processed 16S rDNA amplicon sequencing data, using the command
above and the commands below, twice. In the first pass, we aligned unique
sequences against Silva v. 132 database and proceeded with that alignment.
However, because of the low sequence similarity between some of the planthopper
symbiont sequences and the references, the suboptimal alignment resulted in
some spurious OTUs. Hence, we chose the representative sequences for all OTUs,
aligned them against each other, manually curated the alignment, and used it as
the alignment reference during the second pass.
```

```
######## Getting basic information about the alignment
summary.seqs(fasta=16S.trim.contigs.trim.unique.abund.align)

######## Removing unaligned sequences
screen.seqs(fasta=16S.trim.contigs.trim.unique.abund.align,
count=16S.trim.contigs.trim.abund.count_table, start=1, end=273, minlength=220)

######## Removing any columns containing only gap characters
filter.seqs(fasta=current, vertical=T, trump=.)

######## Re-selecting unique sequences
unique.seqs(fasta=16S.trim.contigs.trim.unique.abund.good.filter.fasta,
count=16S.trim.contigs.trim.abund.good.count_table)

######## Chimera filtering using UCHIME
chimera.uchime(fasta=16S.trim.contigs.trim.unique.abund.good.filter.unique.fast
a, reference=self,
count=16S.trim.contigs.trim.unique.abund.good.filter.count_table,
dereplicate=f, mindiv=0.35, processors=20, minh=0.5, xn=3)

######## Removing chimeric sequences
remove.seqs(accnos=current, fasta=current, count=current)

######## Taxonomic classification of sequences
classify.seqs(fasta=current, count=current,
reference=/mnt/matrix/symbio/db/silva.nr_v132.align,
taxonomy=/mnt/matrix/symbio/db/silva.nr_v132.tax, cutoff=80)

######## Removing sequences not classified as Bacteria
remove.lineage(fasta=current, count=current, taxonomy=current,
taxon=Chloroplast-Mitochondria-Archaea-Eukaryota)

######## Getting sequence classification summary
summary.tax(taxonomy=current, count=current)

######## Changing file names
rename.file(fasta=current, count=current, taxonomy=current, prefix=16S)

######## Computing pairwise distance matrix
dist.seqs(fasta=16S.fasta, processors=24, cutoff=0.05)

######## Clustering OTUs using nearest neighbor method
cluster(column=current, count=16S.count_table, cutoff=0.03, method=nearest)

######## Assigning sequences to OTUs based on 97% identity
bin.seqs(list=current, fasta=current, label=0.03)

######## Creating a 97% OTU table
make.shared(list=current, count=current, label=0.03)
```
